# Aerobic exercise decreases the number and transcript expression of inflammatory M1 macrophages and CD8+ T cells in the epicardial adipose tissue of female pigs

**DOI:** 10.1101/2025.02.02.635562

**Authors:** Irshad Ahmad, Shreyan Gupta, Micah Thomas, James J. Cai, Cristine L. Heaps, Annie E. Newell-Fugate

## Abstract

**Background:** Epicardial adipose tissue (EAT) regulates coronary artery function via lipid metabolism and immune cell recruitment. Increased EAT is a risk factor for coronary artery disease (CAD), but aerobic exercise mitigates CAD. The effect of aerobic exercise on immune cells in EAT is unknown. We hypothesized that aerobic exercise creates an anti-inflammatory environment characterized by increased M2 macrophages and up-regulation of anti-inflammatory cytokine transcripts in EAT.

**Methods:** Female Yucatan pigs (n=7) were allocated to sedentary or exercised groups. To mimic CAD, a coronary artery was chronically occluded or remained non-occluded. EAT samples were processed for bulk and single nuclei transcriptomic sequencing.

**Results:** Sub-clustering identified immune, endothelial, smooth muscle, adipocytes, adipocyte progenitor cells (APSCs), and neuronal cells, with adipocytes and APSCs being dominant. Non-occluded sedentary EAT had the largest percentage of M1 macrophages and CD8+ T cells. Irrespective of occlusion, sedentary EAT had the largest fraction of cells expressing genes in the tumor necrosis factor (TNF) superfamily. Irrespective of occlusion, exercise upregulated peroxisome proliferator-activated receptor (*PPAR*) gamma (*G*) expression and enriched PPAR signaling pathways in adipocytes, macrophages, and T cells. However, *PPARG* expression was lowest in CD8+ T cells from non-occluded exercised EAT. The greatest number of significant cell-cell communications between adipocytes and immune cells via growth factors and adhesion molecules occurred in occluded sedentary EAT.

**Conclusion:** Aerobic exercise mitigates the proinflammatory nature of EAT in CAD via modulation of immune cell subpopulations, decreased TNF superfamily and increased *PPARG* gene expression, and decreased growth factor communication between adipocytes and immune cells.

## Introduction

Epicardial adipose tissue (EAT) mechanically supports the coronary arteries, releases beneficial adipokines, such as adiponectin, and is an energy source for cardiomyocytes, endothelial cells, and smooth muscle cells.^1^ Unlike other adipose tissue depots, EAT lacks distinct boundaries with neighboring tissues and shares its microcirculation with the myocardium.^2^ Due to this close anatomical relationship with the heart, EAT modulates coronary artery function through paracrine secretion of anti- and pro-inflammatory adipokines.^3^ Furthermore, EAT plays a critical role in the local regulation of heart function via lipid metabolism.^4^ During periods of high energy demand, EAT releases free fatty acids to serve as energy substrates for the surrounding tissues.^5^ Within EAT, released free fatty acids can activate the transcription of peroxisome proliferator-activated receptors (PPARs), which stimulates controlled adipogenesis and lipid metabolism.^3, 6^

Although EAT has many beneficial effects on the surrounding tissues, excessive expansion of EAT is a risk factor for coronary artery disease (CAD) due to its increased production of pro-inflammatory molecules, which subsequently invade the adjacent myocardium.^3^ Moreover, as EAT expands and outgrows its vascular supply, hypoxia ensues, which down-regulates anti-inflammatory factors such as adiponectin and causes decreased transcription of antioxidant and thermoregulatory genes such as uncoupling protein 2 (UCP2) and PPAR gamma (PPARG).^7^ PPARG is a master regulator of lipid metabolism in many cell types and is the predominant controller of adipogenesis.^7^ Indeed, myocardial cells can undergo adipogenic differentiation when they express PPARG.^6^ Although PPARG is predominantly expressed in adipocytes, it is also found in low levels in immune cells such as macrophages and T cells.^8^ As EAT is a heterogeneous tissue containing adipocytes, adipocyte progenitor cells, endothelial cells, smooth muscle cells, and various resident and infiltrative immune cells, the role of PPARG in these cell types should be considered.

In response to microenvironmental stimuli, macrophages in EAT exhibit plasticity, altering their polarization between M1 pro-inflammatory and M2 anti-inflammatory phenotypes.^9^ This phenotypic shift occurs in response to local cytokines and inflammatory substances, with M1 polarization activated by interferon-γ (IFN-γ) or lipopolysaccharide (LPS) and M2 polarization activated by interleukin (IL)-13 or IL-4.^10^ PPARG is responsible for the differentiation of monocytes to the M2 anti-inflammatory macrophage phenotype^11^ and promotes the uptake of oxidized low-density lipoprotein (LDL), which contributes to foam cell formation.^12^ Thus, PPARG in EAT may play a significant role as a potential mediator against CAD.^13^

EAT volume contracts in response to aerobic exercise^14^ due to an increased lipolytic rate of the tissue.^15^ Moreover, most studies have demonstrated a correlation between decreased EAT volume and changes in circulating markers of cardiometabolic health, such as decreased C-reactive protein (CRP), LDL, total cholesterol^16^, and TNF-α, and increased adiponectin.^17^ However, the effect of aerobic exercise on immune cells and genes related to metabolism in EAT is unknown. In this study, we examined the impact of aerobic exercise on the single nuclei transcriptome in EAT from sexually mature female Yucatan miniature swine with experimentally induced chronic ischemic heart disease.

## Materials and Methods

The analytic methods and data are available by request to other researchers to reproduce the results or replicate the procedures.

### Experimental animal procedures

Animal protocols were approved by the Texas A&M University Institutional Animal Care and Use Committee and conformed to the National Institutes of Health (NIH) Guide for Care and Use of Laboratory Animals, 8^th^ edition, revised 2011. Adult female Yucatan miniature swine (6-7 months of age) were surgically instrumented with an ameroid constrictor around the proximal left circumflex coronary artery as previously described.^18–20^ The ameroid constrictor progressively narrows the artery and induces an inflammatory response resulting in gradual occlusion, with total obstruction occurring at approximately two to three weeks post-operatively.^21^ Pigs recovered for eight weeks post-operatively before the sedentary or exercise training experimental regimen was initiated.

### Sedentary and exercise protocols

Exercise-trained female pigs underwent a progressive treadmill exercise training program, five days/week for 14 weeks, as described previously.^20, 22^ Briefly, speed and duration of the exercise training sessions were progressively increased so that during the last three weeks of training, animals ran at 4-5.5 mph for 60 minutes and at 6 mph for 5-15 minutes. Grade of the treadmill was maintained at 0% throughout the exercise bouts. Exercise-trained and sedentary swine were fed once daily immediately after the exercise training session as positive reinforcement for the exercise bout, with water provided *ad libitum*. Sedentary pigs maintained normal activity in their pens throughout the experimental protocol. Effectiveness of the exercise training regimen for these animals was determined by comparing heart-to-body weight ratio and skeletal muscle citrate synthase activity. Data on these parameters for the study animals was presented in Ahmad et. al. 2024.^20^

### Euthanasia for tissue collection

Pigs were anesthetized with intramuscular administration of xylazine (2.25 mg·kg^-1^) and ketamine (35 mg·kg^-1^), fitted with an anesthesia mask, induced with 3% isoflurane and then intubated and a surgical plane of anesthesia was maintained with 3% isoflurane and auxiliary O_2_. A left lateral thoracotomy was performed in the fourth intercostal space, followed by administration of heparin (500 U·kg^-1^, i.v.). Hearts were removed, placed in Krebs bicarbonate buffer (0-4 °C) and weighed. Krebs bicarbonate buffer contained (in mM): 131.5 NaCl, 5 KCl, 1.2 NaH2PO4, 1.2 MgCl2, 2.5 CaCl2, 11.2 glucose, 13.5 NaHCO3 and 0.025 EDTA.

### Isolation of coronary epicardial adipose tissue

Following humane euthanasia, epicardial perivascular fat was sectioned from both the non-occluded left anterior descending coronary artery and the collateral-dependent left circumflex coronary artery. Perivascular fat was snap-frozen in liquid nitrogen and stored at -80 °C for later analysis. Subsequent visual inspection of the ameroid constrictor during dissection of the left circumflex artery under a dissection microscope during dissection of the left circumflex artery indicated 100% occlusion in all pigs used in this study.

### Bulk RNA isolation and sequencing

EAT (occluded and non-occluded tissue pooled) from exercise-trained (n=4) and sedentary (n=3) female pigs had total RNA extracted with TRIzol® (Thermofisher Scientific, Waltham, MA). Total RNA was quantified (Nanodrop 3300, Thermofisher, Wilmington, DE) followed by bioanalysis (Agilent 2100 Bioanalyzer, Agilent Technologies, Inc., Santa Clara, CA) for RNA quantity and quality. RNA integrity scores fell within the following range: 6.2–9.1. cDNA libraries were prepared, and sequencing was performed at the Molecular Genomics Core at Texas A&M University.

### Single-nucleus RNA isolation and sequencing

Isolation of EAT nuclei (occluded and non-occluded isolated separately) was performed following the “Daughter of Frankenstein protocol for nuclei isolation from fresh and frozen tissues using OptiPrep continuous gradient V.2”^23^ and as previously described.^20^ Nuclei were resuspended in 0.1% BSA in PBS and immediately processed to generate single-nuclei RNA libraries using the microdroplet-based RNA. Nuclei samples were diluted, if necessary, to a target concentration of between 500 – 1,500 nuclei/µL and used for single nuclei RNA sequencing library preparation.^24^

Single nuclei 3’ sequencing libraries were generated following the 10x Genomics dual index manual preparation v3.1 reagent kit with a target nuclei recovery of approximately 5,000 – 10,000 nuclei for each reaction. GEM partitioning was carried out in the 10x Genomics Chromium X system and following library construction following the manufacturer’s protocol. Sequencing libraries were quantified with the Qubit 4.0 Fluorometer high sensitivity dsDNA detection kit (Thermofisher). Library sizes were assessed using the Agilent TapeStation 4200, D1000 DNA tape system (Agilent Technologies). All sequencing library concentrations were normalized to 4 nM and pooled at equimolar ratios for sequencing on an Illumina NovaSeq 6000 2x150 sequencing run to generate a minimum of 200 million, paired-end sequencing reads for each sample. Sequencing reads were loaded into the 10x Genomics Cell Ranger software for sequence alignment, filtering, barcode and unique molecular identifier (UMI) counting.

### Bioinformatic Analysis

#### Bulk RNA Analysis

Raw bulk RNA sequencing data underwent quality control using FastQC (v0.11.9) (http://www.bioinformatics.babraham.ac.uk/projects/fastqc). Read alignment to the reference genome was performed with Bowtie2 (v2.3.5.1)^25^ using default parameters. Gene-level counts were obtained with htseq-count (v2.0.2).^26^ Data normalization was carried out using the upper-quartile method to account for library size differences.

Differential expression analysis was performed in female pigs, comparing the exercised and sedentary groups using R (v4.1.0). The edgeR package (v3.34.1)^27^ was used for count filtering and dispersion estimation, and limma (v3.48.3)^28^ was used with the voom transformation for linear modeling. A total of 13,663 genes were included in the analysis, identifying 495 genes (275 upregulated and 220 downregulated) as differentially expressed genes (DEGs) (P-value < 0.05 and absolute Log2 fold-change (log2FC) > 1.5). DEGs were plotted as a volcano plot.

Log2FC values for all genes were imported into the ‘fgsea’ R package^29^ for gene set enrichment analysis (GSEA) against the KEGG Human 2023 gene sets. Pathways with an absolute normalized enrichment score (NES) > 1.645 and adjusted P-value < 0.05 were significantly enriched. A total of 19 pathways met these criteria.

#### Single nuclei RNA analysis

UMI count matrices for each single-cell sample were generated by aligning reads to the genome (Landrace_pig_v1 (GCA_001700215.1) using 10X Genomics Cell Ranger software. Data was analyzed by exercise status alone (Exercise (Ex) versus Sedentary (Sed)) or the combination of exercise with occlusion status (N_Ex: non-occluded-exercised, O_Ex: occluded-exercised, N_Sed: non-occluded sedentary, O_Sed: occluded-sedentary). The count matrices for all four treatment groups were merged into one Seurat object without batch correction using the Seurat v4 R package.^30^ Cells with less than 500 feature counts were filtered out. Genes expressed in less than 15 cells also were filtered out. The ‘*NormalizeData’* function was used for normalization. This function divided the gene expression counts for each cell by the total counts for that cell and multiplied the value by 10,000. Then, the counts were Log1p transformed and scaled using the ‘*ScaleData*’ function. ‘*FindVariableFeatures’* function was used to identify the top 2,000 highly variable genes. These 2,000 genes were then used for Principal Component Analysis (PCA). Uniform Manifold Approximation and Projection (UMAP) was performed using the first 50 principal components to project the data onto a three-dimensional (3D) space for visualization. Clustering was performed using the K-means algorithm on the 3D UMAP projection to identify 50 cell clusters. The K-means algorithm assigns data points (cells) to the nearest centroids (cluster center), and recomputes the centroids based on the predefined number of clusters.^31^ After identifying cell clusters, immune cells were separated from the other cell clusters, and their UMAP projections were recalculated. After parsing out the immune cell cluster, its cell constituents were clustered into four groups using the K-means algorithm and the UMAP projections. Significant marker genes for each cluster were identified by performing differential analysis using the Wilcoxon rank sum test. Genes with minimum percent expressed in cluster > 25%, |log2(FC)| > 0.25, and false discovery rate (FDR) < 0.05 were considered marker genes. Each cluster was annotated using the top 10 computed marker genes and canonical cell type markers from PanglaoDB^32^ and the pig atlas (https://dreamapp.biomed.au.dk/pigatlas/).

Differential expression analysis was performed using the Wilcoxon rank-sum test for each cell type and across all four treatment groups. Significantly differentially expressed genes (DEG) were selected by a FDR cut-off of 0.05 and |log2(FC)| >2. DEG for each cell type were used for gene set enrichment analysis with the Kyoto Encyclopedia of Genes and Genomes (KEGG) pathways database using fgsea R package.^33^ Cell scores for gene sets were calculated using the AUCell ^34^ for gene sets. Gene sets were obtained from the KEGG Human Pathway Database. Visualization was carried out using Seurat’s inbuilt plotting function and the ggplot R library.

The CellChat (v1) R package^35^ was used to infer cell-cell communication within and across cell types in the single nuclei dsata set. To quantitatively analyze the cell-cell interaction, all cells from the four treatment groups (N_Ex, N_Sed, O_Ex, O_Sed) were pooled and grouped by cell types for CellChat analysis. Default parameters were used. The human ligand-receptor database, without any selection, was used. For each cell type, significantly differentially overexpressed ligand-receptor pairs were identified. Next, the communication likelihood was computed using the average expression values of a ligand in one cell type compared to that of a receptor or cofactor in another cell type. Significant interactions were pruned by permutation tests (P value < 0.05). An intercellular communication network was generated along with communication probabilities that measure the strength of relationships between each ligand-receptor pair. The communication network for each signaling pathway was generated by summing up all corresponding ligand-receptor interactions for that pathway. The CellChatDB was subdivided to identify paracrine, endocrine, and juxtacrine signaling pathways. The visualization was carried out using CellchatR’s inbuilt plotting functions.

## Results

### Aerobic exercise enriches pathways related to immune function and cytoskeletal organization and negatively enriches pathways related to nutrient utilization and metabolism in EAT

There were 13,663 DEG, of which 338 were significantly up-regulated, and 293 were significantly down-regulated in the exercise-trained group (**Figure 1A**). The most significantly upregulated genes in EAT from exercise-trained pigs were: shisa family member 2 (*SHISA2*), cadherin 7(*CDH7*), heparin-binding EGF-like growth factor (*HB-EGF*), family with sequence similarity 71 member E1 (*FAM71E1*), and pleckstrin homology-like domain family A member 2 (*PHLDA2*) (**Figure 1A**; **Supplemental Dataset 1**). Genes that were most significantly down-regulated in EAT from exercise-trained pigs were: cytochrome P450 family 2 subfamily B member 6 (*CYP2B6*), acetylcholinesterase (*ACHE*), polypeptide N-acetyl-galactos-aminyl-transferase like 5 (*GALNTL5*), Metallothionein 2A *(MT2A*), claudin-4 (*CLDN4*), estrogen-related receptor gamma (*ESRRG*), keratin 14 (*KRT14*), galactose-3-O-sulfotransferase 3 (*GAL3ST3*) (**Figure 1A**; **Supplemental Dataset 1**)

**Figure 1.**
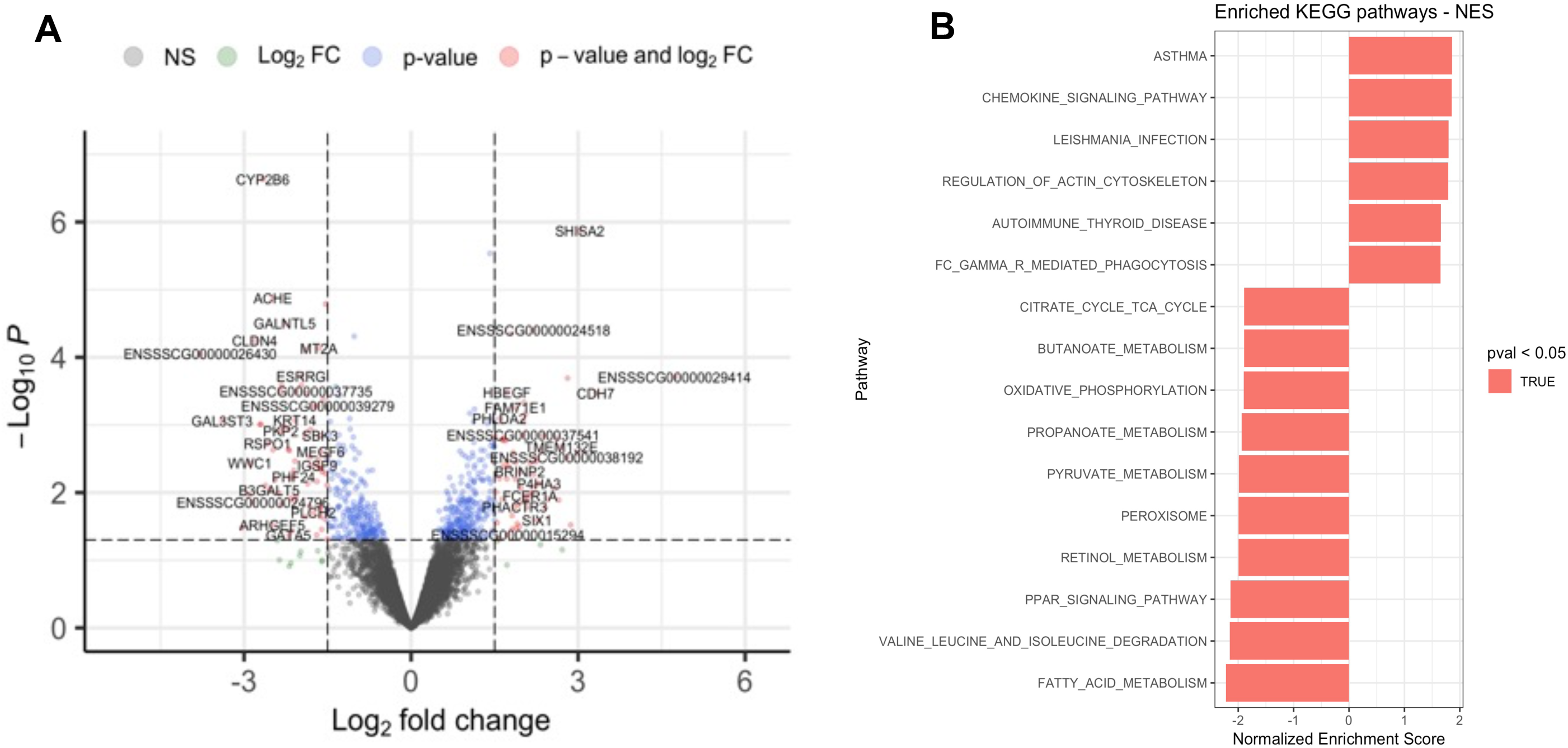
Analysis of differentially expressed genes and enrichment of KEGG pathways in the EAT from exercised-trained female pigs. (A) Volcano plot of up-regulated and down-regulated genes in exercise-trained versus sedentary pigs. (B) Identification of KEGG pathways significantly enriched (upregulated genes) or negatively enriched (downregulated genes) in exercise-trained pigs. The threshold set points for DEG were fold change > 2 and adjusted P value < 0.05. NS: not significant. Log_2_FC: log base 2-fold change. Log_10_: log base 10

Significant enrichment based on KEGG pathway analysis was found for asthma, chemokine signaling pathway, leishmania infection, regulation of actin cytoskeleton, autoimmune thyroid disease, and FcγR-mediated phagocytosis in exercise-trained pigs (**Figure 1B**). In contrast, pathways with negative enrichment scores in exercise-trained pigs were: TCA cycle, butanoate metabolism, oxidative phosphorylation, propanoate metabolism, pyruvate metabolism, peroxisome, retinol metabolism, PPAR signaling, valine-leucine-isoleucine degradation, and fatty acid metabolism (**Figure 1B**).

### Non-occluded EAT from sedentary pigs has large numbers of macrophages and T cells

The total number of nuclei derived from single nuclei processing in occluded and nonoccluded exercised and sedentary EAT was 28,473. After removing nuclei with less than 500 feature counts, 27,828 single nuclei were available for downstream analysis. K-clustering analysis and marker gene annotation revealed seven cell populations in the four treatment groups (**Figure 2A-B, Figure S1, Supplemental Dataset 2**). Each cell type showed a specific set of enriched gene expression: adipocytes - perilipin 1 (*PLIN1*) and adiponectin, C1Q, and collagen domain containing (*ADIPOQ*); adipose progenitor and stem cells (APSC) – platelet-derived growth factor receptor alpha (*PDGFRA*) and collagen type VI alpha 1 chain (*COL6A1*); endothelial cells - von Willebrand factor (*VWF*) and fatty acid binding protein 5 *(FABP5*); macrophages - CD163 molecule (*CD163*) and stabilin 1 *(STAB1);* neurons - synuclein alpha (*SNCA*), neurexin 1 *(NRXN1*); smooth muscle cells - actin alpha 2, smooth muscle (*ACTA2*) and gap junction protein gamma (*GJC1*); T cells - Neurocalcin delta (*NCALD) and* Src kinase-associated phosphoprotein 1(*SKAP1)* (**Figure 2C**; **Supplemental Dataset 2; Table S1**). Although APSCs are characterized by upregulated *PDGFRA* and *COL6A1* gene expression, these genes have increased expression in only 20-30% of APSCs. Nearly all other marker genes are expressed in > 50% of each cell type except for *ACTA2,* which is only expressed in 30% of smooth muscle cells (**Figure 2C**).

**Figure 2.**
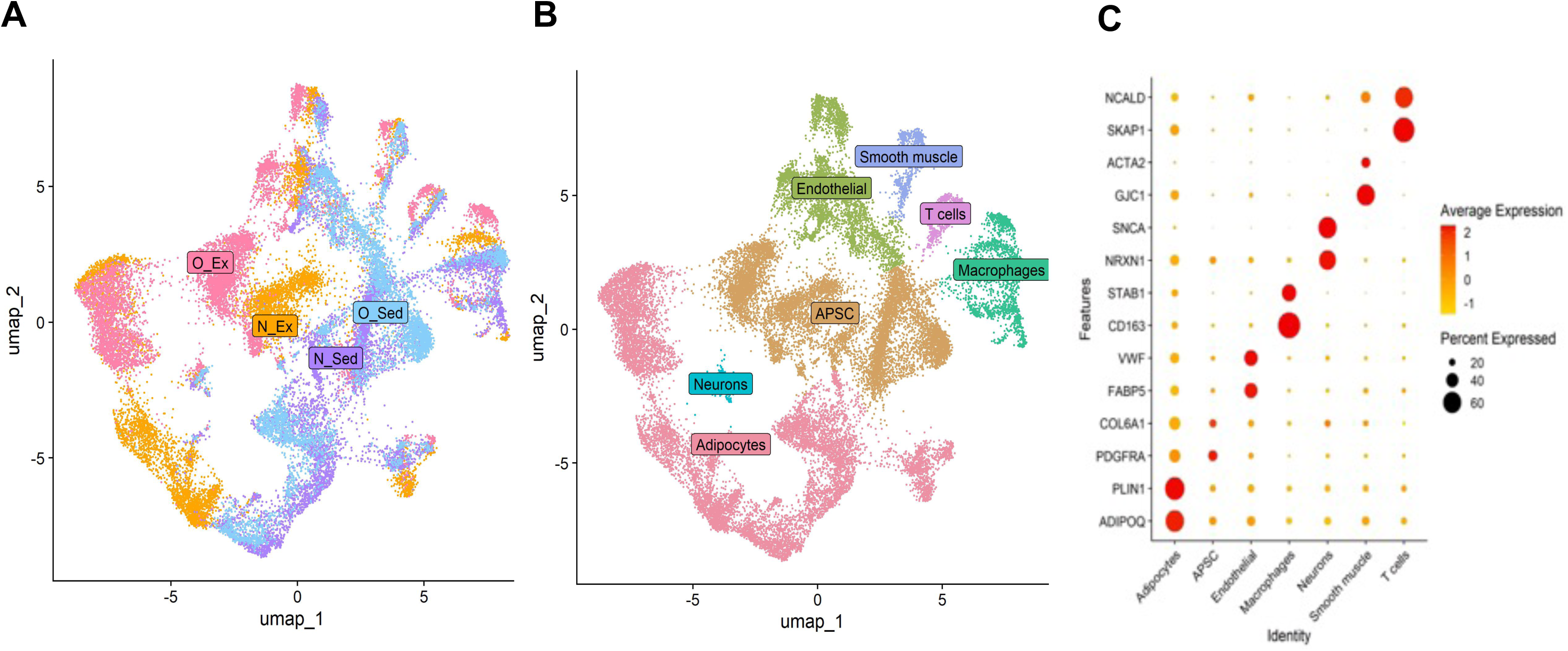
Distribution by treatment group and cell type of nuclei from occluded and non-occluded EAT of exercise-trained and sedentary female pigs. (A) UMAP plot of single nuclei from EAT of exercise-trained and sedentary female pigs showing the origins of nuclei in different treatment groups. (B) UMAP plot of single nuclei from EAT of female pigs showing seven major cell types from sedentary (occluded and nonoccluded) and exercised (occluded and non-occluded) tissue samples. (C) Bubble heatmap demonstrating expression levels of marker genes for each cell (nuclei) type. N: non-occluded; O: occluded; Ex: exercise trained; Sed: sedentary. APSC: adipocyte progenitor cells.

Irrespective of exercise status, non-occluded EAT had a large number of adipocytes (3437 cells in N_Ex, 2985 cells in N_Sed) (**Figure 3A-B**). By contrast, APSCs were most abundant in N_Sed and O_Ex EAT. Endothelial and smooth muscle cell numbers in EAT were similar across treatment groups except for the N_Ex, which had lower numbers of these cell types (**Table S2**). Neuron abundance in EAT was the same across all treatment groups. N_Sed had a higher number of T cells (254 cells) and macrophages (1199 cells) compared to any other treatment group (**Figure 3A-B**, **Table S2**). Cell numbers showed batchwise distribution and abundance differences in adipocytes and APSCs in N_Ex and O_Ex EAT. N_Sed showed higher numbers but no difference in batchwise distribution of immune cells and adipocytes (**Figure 3C, Table S2**).

**Figure 3.**
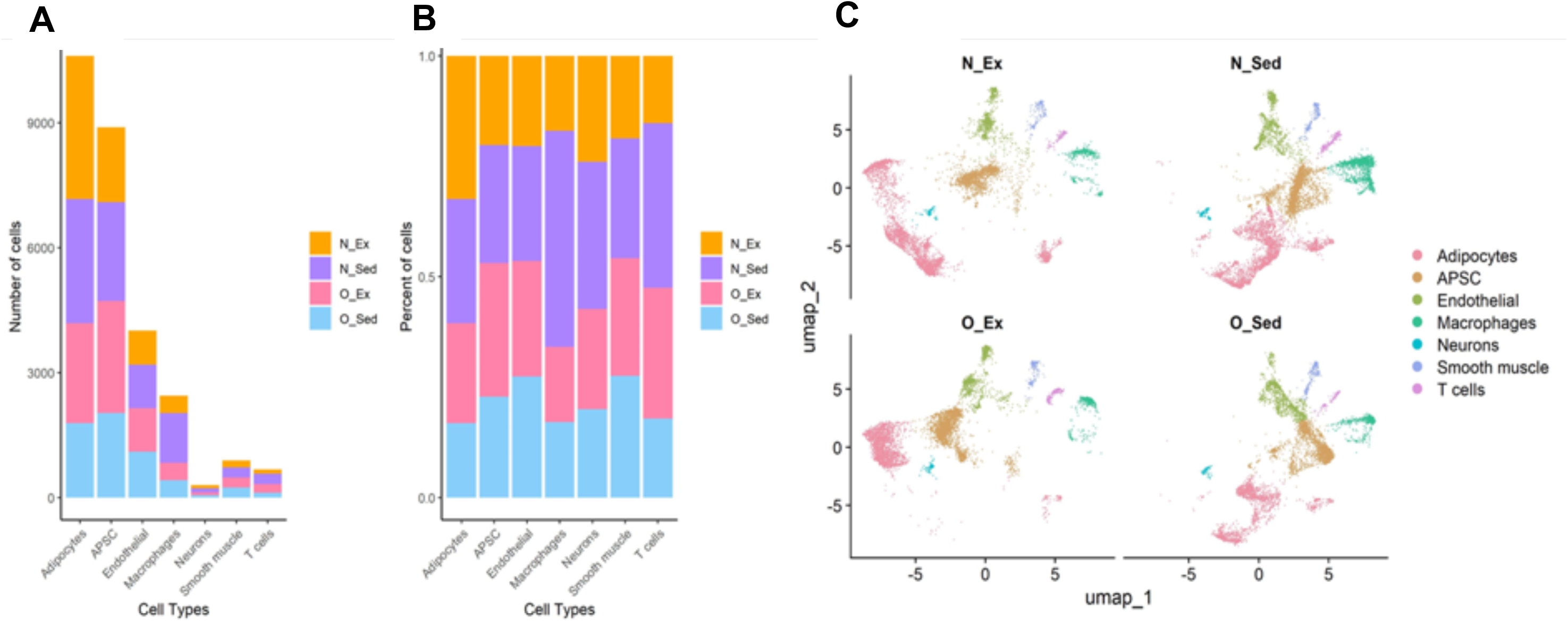
Quantity and distribution of cell types across all treatment groups. (A) Barplot of the number of cell types in each treatment group. (B) Barplot of the percentage of cell types in each treatment group. (C) Batchwise distribution of each cell type. N: non-occluded; O: occluded; Ex: exercise trained; Sed: sedentary. APSC: adipocyte progenitor cells.

### Irrespective of coronary artery occlusion status, aerobic exercise upregulates genes in most cell types in EAT related to unsaturated fatty acid metabolism and PPAR signaling

The differentially expressed genes (DEGs) for adipocytes, macrophages, T cells, endothelial cells, and smooth muscle cells were identified in occluded and non-occluded EAT from exercise-trained and sedentary pigs (**Supplemental Datasets 3 & 4**). Irrespective of treatment group, adipocytes had the greatest number and percentage of DEGs in response to exercise (non-occluded: 677/8445; 8% of genes; occluded: 732/8803; 8.3% of genes; **Figure S2 & S5**). Malic enzyme 1 (*ME1*) and stearoyl-CoA desaturase (*SCD*) were significantly upregulated in response to exercise in adipocytes from both occluded and non-occluded EAT **(Figure S2 & S5**). In N_Ex EAT, genes such as pyruvate dehydrogenase kinase 4 (*PDK4*), fibronectin 1 (*FN1*), and glutathione peroxidase (*GPX3*) were significantly downregulated in adipocytes (**Figure S2**). In contrast, in O_Ex genes such as cysteine-rich transmembrane BMP regulator 1 (*CRIM1*), glycoprotein M6A (*GPM6A*), and PKHD1 like 1 (*PKHD1L1*) were significantly down-regulated in adipocytes (**Figure S5**). In adipocytes from occluded and non-occluded EAT PPAR signaling, retinol metabolism, biosynthesis of unsaturated fatty acids, and pyruvate metabolism were significantly enriched in response to exercise training. In exercised pigs, non-occluded EAT also had significant enrichment of the insulin signaling and adipocytokine and glycolysis-gluconeogenesis pathways. In contrast, occluded EAT from exercised pigs had an enrichment of steroid and terpenoid biosynthetic pathways. Exercise training enriched PPAR signaling in all cell types examined except smooth muscle cells.

With respect to immune cells in EAT, irrespective of occlusion status, acetyl-CoA carboxylase alpha (*ACACA*), ELOVL fatty acid elongase 6 (*ELOVL6*), and *ME1* were significantly upregulated in macrophages, and pyruvate metabolism and biosynthesis of unsaturated fatty acids pathways were enriched in response to exercise training **(Figure S3 & S6**). Similar to adipocytes from non-occluded EAT, macrophages from non-occluded EAT had enrichment of the insulin signaling pathway in response to exercise training (**Figure S3**). On the other hand, in T cells, the occlusion status of EAT significantly affects the types of genes and pathways impacted by exercise training. In T cells from N_Ex EAT, the top genes significantly upregulated were ATP citrate lyase (*ACLY*), *SCD*, and Rho GTPase activating protein 28 (*ARHGAP28)* (**Figure S3**). In T cells from O_Ex EAT, the top genes that were significantly upregulated were *ELOVL6*, *ACACA,* and *ME1* (**Figure S6**). The only significantly enriched pathway in T cells from N_Ex EAT was PPAR signaling; this finding is likely due, in part, to the fact that only 13 genes were significantly upregulated in T cells from N_Ex EAT (**Figure S3**). The enriched pathways in T cells from O_Ex EAT were like those for adipocytes: PPAR signaling, pyruvate metabolism, and biosynthesis of unsaturated fatty acids. Interestingly, pathways significantly negatively enriched in T cells from O_Ex included those related to T cell signaling and metabolism: T cell receptor signaling pathway, Jak-Stat signaling pathway, and natural killer cell-mediated cytotoxicity (**Figure S6**).

In cells that likely comprise small capillaries in EAT – endothelial and smooth muscle cells – genes related to metabolism *ELOVL6*, *ME1*, *ACLY*, and *ACACA* were upregulated (**Figures S4 & S7**). Like adipocytes from O_Ex EAT, endothelial and smooth muscle cells from O_Ex EAT, had downregulation of *GPM6A* and *PKHD1L1* (**Figure S7**). Irrespective of occlusion status, exercise training did not significantly enrich pathways from smooth muscle cells. Similar to adipocytes and irrespective of occlusion status, exercise training significantly enriched PPAR signaling, pyruvate metabolism and biosynthesis of unsaturated fatty acids in endothelial cells **(Figures S4 & S7)**. Furthermore, like adipocytes from N_Ex EAT, endothelial cells from N_Ex EAT had significant enrichment of the insulin signaling pathway (**Figure S4**). In endothelial cells from O_Ex EAT, insulin signaling was also enriched, as was the ribosome pathway and glycerolipid metabolism (**Figure S7**).

### Sedentary EAT has increased numbers of CD8+ T cells expressing TNF superfamily genes

The pro-inflammatory factors synthesized by immune cells act as key mediators in CAD^36^. Therefore, we performed a detailed bioinformatic analysis of the macrophages and T cells in our samples to ascertain the relationship between chronic coronary artery ischemia, exercise training, and immune cells in EAT. N_Sed EAT had the largest number of macrophages (**Figure 4A**). On the other hand, T cells were abundant in EAT from both the N_Sed and O_Ex groups (**Figure 4A-B**). Macrophages had a higher expression of *CD163* and *STAB1* than T cells did. On the other hand, T cells had a higher expression of *SKAP, NCALD, and TOX* (**Figure 4C**). T cell and macrophage clustering analysis demonstrated a separation of both the T cell and macrophage populations from O_Ex EAT from these cell populations from the rest of the treatment groups (**Figure 4D-E**). Based on the degree of separation, the O_Ex EAT macrophages differed more from all other macrophages than the O_Ex EAT T cells differed from all other T cells. Macrophages from sedentary EAT were similar irrespective of occlusion status.

**Figure 4.**
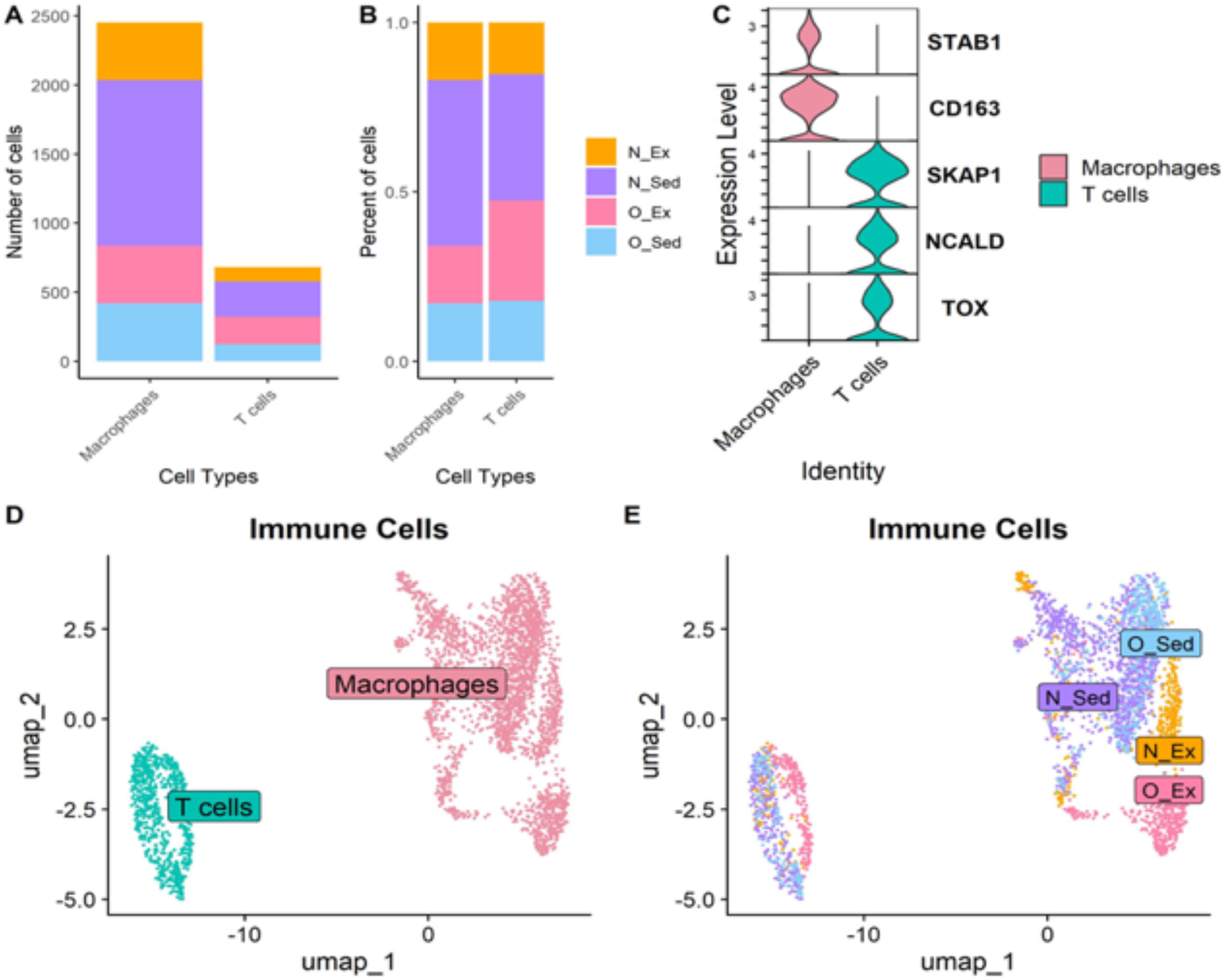
Characterization of the cell number and gene signatures of macrophages and T cells in EAT in response to exercise and coronary artery occlusion. (A) Barplot the number of cells in each treatment group. (B) Barplot the percentage of cells in each treatment group. (C) Violin plot of the expression of selected marker genes in each cell type from EAT. (D) UMAP plot of T cell and macrophage clusters identified by unsupervised clustering. (E) UMAP plot of single nuclei from T cells and macrophages showing the origins of nuclei in different treatment groups. N: non-occluded; O: occluded; Ex: exercise trained; Sed: sedentary. STAB1: stabilin 1. CD163: CD163 molecule. SKAP1: Src kinase-associated phosphoprotein 1. NCALD: neurocalcin delta. TOX: thymocyte selection associated high mobility group box.

Further analysis of the T cell population allowed partitioning of the CD8+ T cells, the most abundant T cell type in the EAT samples (**Figure 5; Figure S8**). Irrespective of occlusion status, EAT from sedentary pigs had more T cells than EAT from exercised pigs (**Figure 5A**). However, O_Ex EAT had double the number of T cells as N_Ex EAT. The CD8+ cell population was classified by increased expression of the following gene markers: CD8 subunit alpha (*CD8A),* C-C motif chemokine ligand 5 (*CCL5),* CD2 molecule *(CD2),* granzyme H *(GZMH),* and transforming growth factor beta receptor 2 (*TGFBR2*) (**Figure 5B**). CD8+ T cells from O_Sed EAT had the highest expression levels of all CD8+ T cell gene markers compared to EAT from all other treatment groups. *CD8A* and *TGFBR2* were highly expressed in CD8+ T cells from all treatment groups. CD8+ T cells from N_Ex EAT also had high expression of CD2. Irrespective of occlusion status, sedentary EAT had the highest expression of tumor necrosis factor (TNF) superfamily gene expression (**Figure 5C**). Within exercise treatment group, non-occluded EAT had higher expression of TNF superfamily genes than occluded EAT; however, this finding only reached significance for the occluded EAT (**Figure 5C**). The *PPARG* expression level was similar across treatment groups when analyzed with all T cells batched together (**Figure S9**). However, when analyzing CD8+ T cells only, the PPARG expression level was very low in the N_Ex treatment group as compared to all other treatment groups (**Figure S9**).

**Figure 5.**
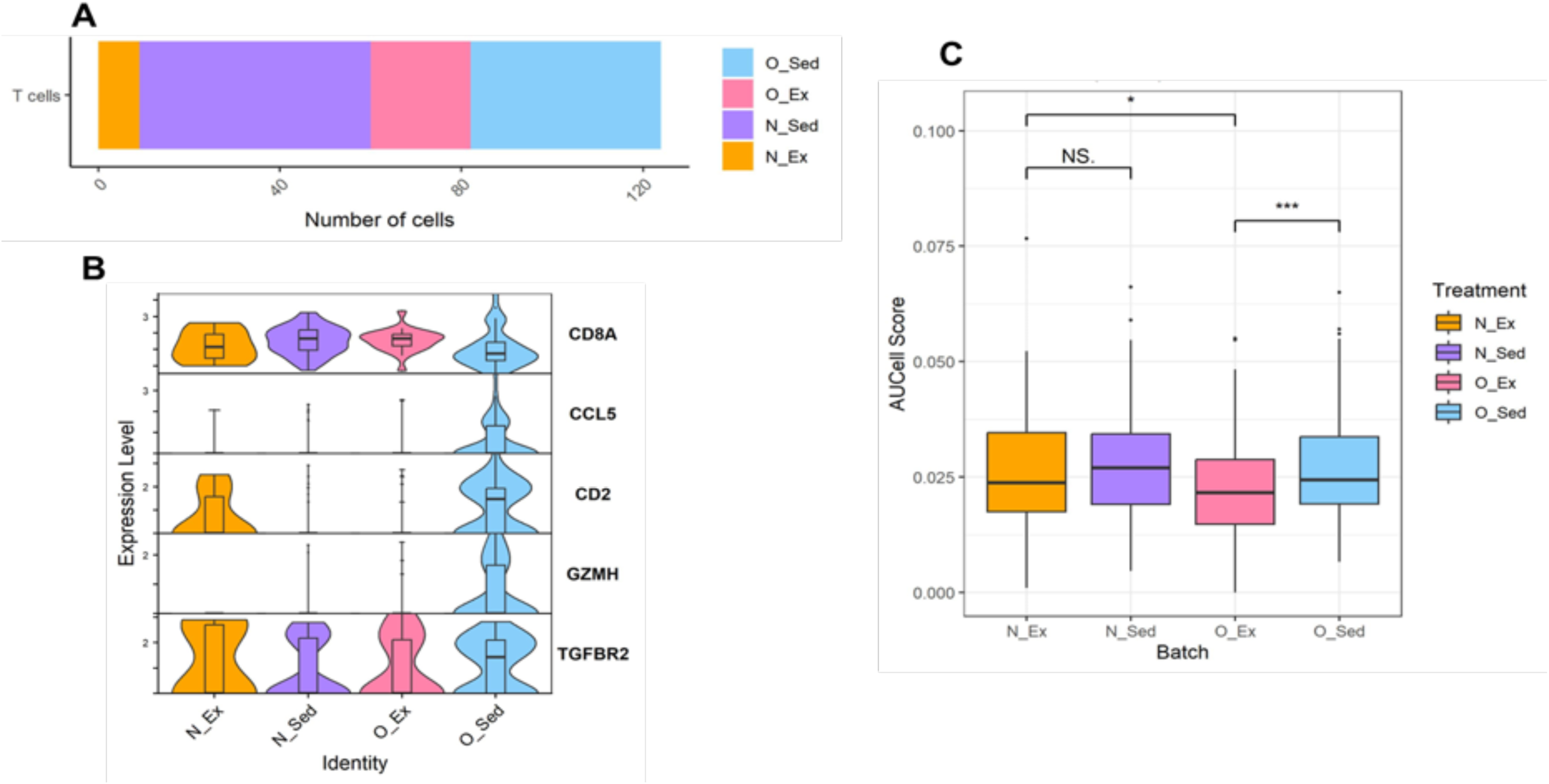
Number and gene signature of CD8+ T cells in EAT by exercise and coronary artery occlusion treatment group. (A) Barplot of the CD8+ T cell number in EAT from each treatment group. (B) Violin plot of the expression of selected marker genes for identification of the CD8+ T cell population in EAT from each treatment group. (C) Box plot of AUCell scores of T cells from EAT of each treatment group expressing genes in the tumor necrosis factor (TNF) superfamily. N: non-occluded; O: occluded; Ex: exercise trained; Sed: sedentary. CD8A: CD8A molecule. CCL5: C-C motif chemokine ligand 5. CD2: CD2 molecule. GZMH: granzyme H. TGFBR2: transforming growth factor receptor 2. AUCell uses the Area Under the Curve to determine if a subset of the input gene set is enriched in the expressed genes for a given cell type. NS: not significant. * P < 0.05. *** P < 0.001.

### Sedentary EAT contains an increased number of pro-inflammatory macrophages M1 expressing TNF superfamily genes

Based on their polarization, macrophages can be broadly categorized into M0, M1, and M2 phenotypes. M0 macrophages are naïve, inactivated cells. M1 macrophages participate in pro-inflammatory responses, while M2 macrophages exhibit an anti-inflammatory profile^37^.

M0 macrophages are phenotypically different from M1 and M2 macrophages (**Figure 6A**). A subpopulation of M2 macrophages separated from the rest of the M2 macrophages was more similar to M0 macrophages than other M2 macrophages (**Figure 6A)**. N_Sed EAT had the most macrophages of all the treatment groups, with M0 and M1 macrophages being the most abundant (**Figure 6B)**. Irrespective of occlusion status, EAT from sedentary pigs had a higher percentage of M1 macrophages than EAT from exercised pigs (**Figure 6B**). EAT from exercised pigs had a higher percentage of M0 macrophages than EAT from sedentary pigs. The percentage of macrophages that were M2 polarization was similar between the four treatment groups **(Figure 6B)**. M1 macrophages had the highest expression level of CD80 molecule (*CD80*) and interferon regulatory factor 5 (*IRF5*). In contrast, M2 macrophages had the highest expression level of CD209 molecule *(CD209),* and transforming growth factor beta 1 *(TGFB1)* (**Figure 6C**). Irrespective of occlusion status, sedentary EAT had the highest expression of TNF superfamily genes (**Figure 6D**).

**Figure 6.**
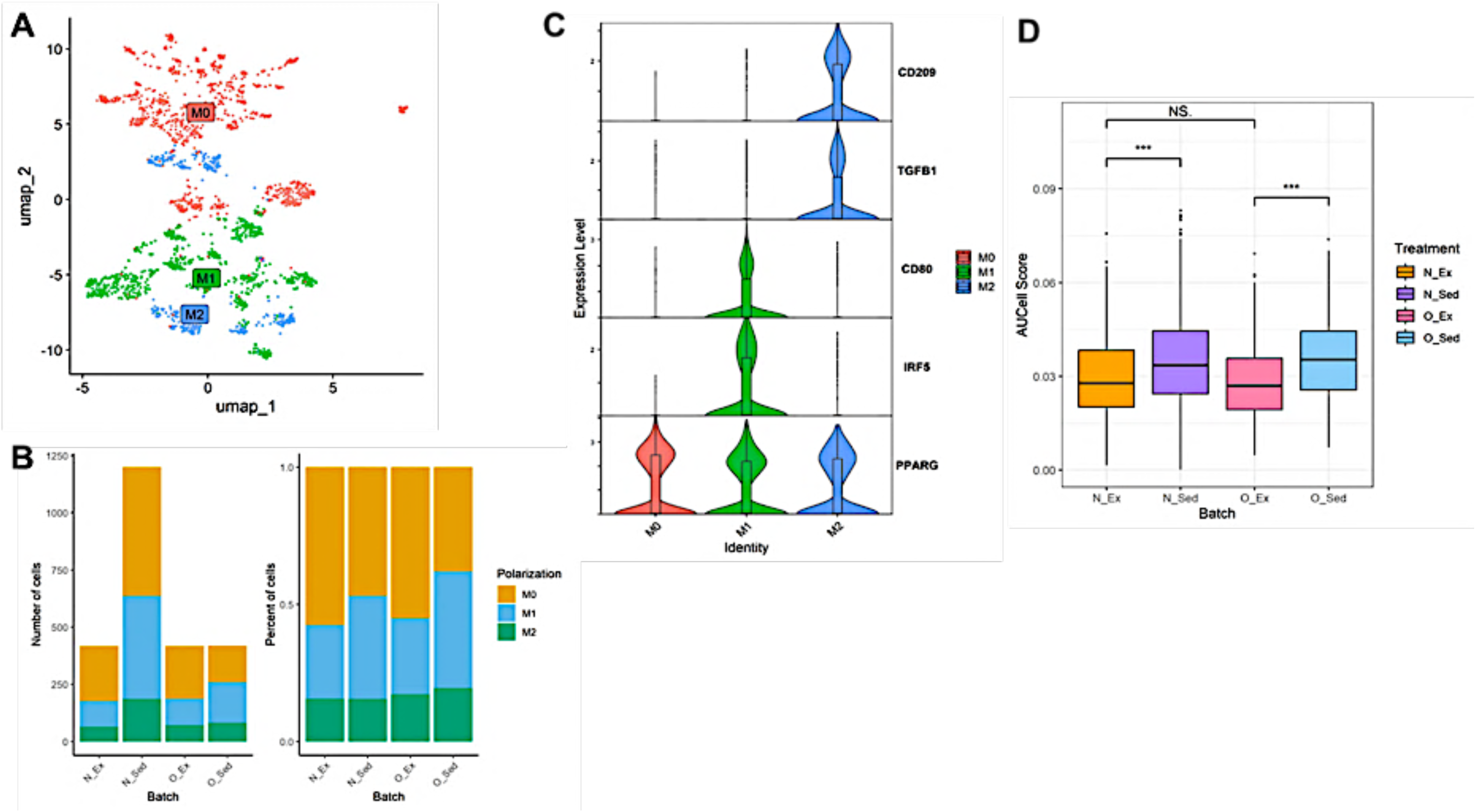
Functional analysis of macrophage populations in EAT in response to exercise and coronary artery occlusion status. (A) UMAP plot of macrophages classified by respective marker genes. (B) Barplot of the number and percentage of M0, M1, and M2 macrophages in each treatment group. (C) Violin plot showing the expression level of selected marker genes in each macrophage class. (D) Box plot of AUCell scores of macrophages from EAT of each treatment group expressing genes in the tumor necrosis factor (TNF) superfamily. N: non-occluded; O: occluded; Ex: exercise trained; Sed: sedentary. CD209: CD209 molecule. TGFB1: transforming growth factor beta 1. CD80: CD80 molecule. IRF5: interferon regulatory factor 5. PPARG: peroxisome proliferator-activated receptor gamma

*PPARG* expression was similar across all macrophage subpopulations (**Figure 6C**). Genes related to PPARG signaling were upregulated in macrophages in EAT from O_Ex pigs (**Figure S10**). In contrast, occlusion status did not affect *PPARG* or lipoprotein lipase (*LPL*) expression in adipocytes from EAT, but exercise status did (**Figure S10**). On the other hand, exercise did not affect the expression of peroxisome proliferator-activated receptor delta (*PPARD)*, *SCD,* or *ME1* in adipocytes from EAT, but occlusion status did (**Figure S10**).

### Adipocytes, adipocyte stem cells, and immune cells from occluded sedentary EAT communicate via growth factors and cell adhesion molecules

CellChat found 393 ligand-receptor pairs within the macrophages, T cells, adipocytes, and APSC from EAT. A weighted communication-directed plot based on the significant connections between cell types found that adipocytes and APSCs engaged in the most substantial interactions with the immune cells (**Figure 7A**). Adipocytes had the greatest number of communications, most of which were with APSCs, followed by macrophages. O_Ex EAT had the greatest number of incoming and outgoing interactions across all cell types (**Figure S11**). However, N_Sed EAT had the greatest number of incoming interactions in adipocytes (**Figure S11).**

**Figure 7.**
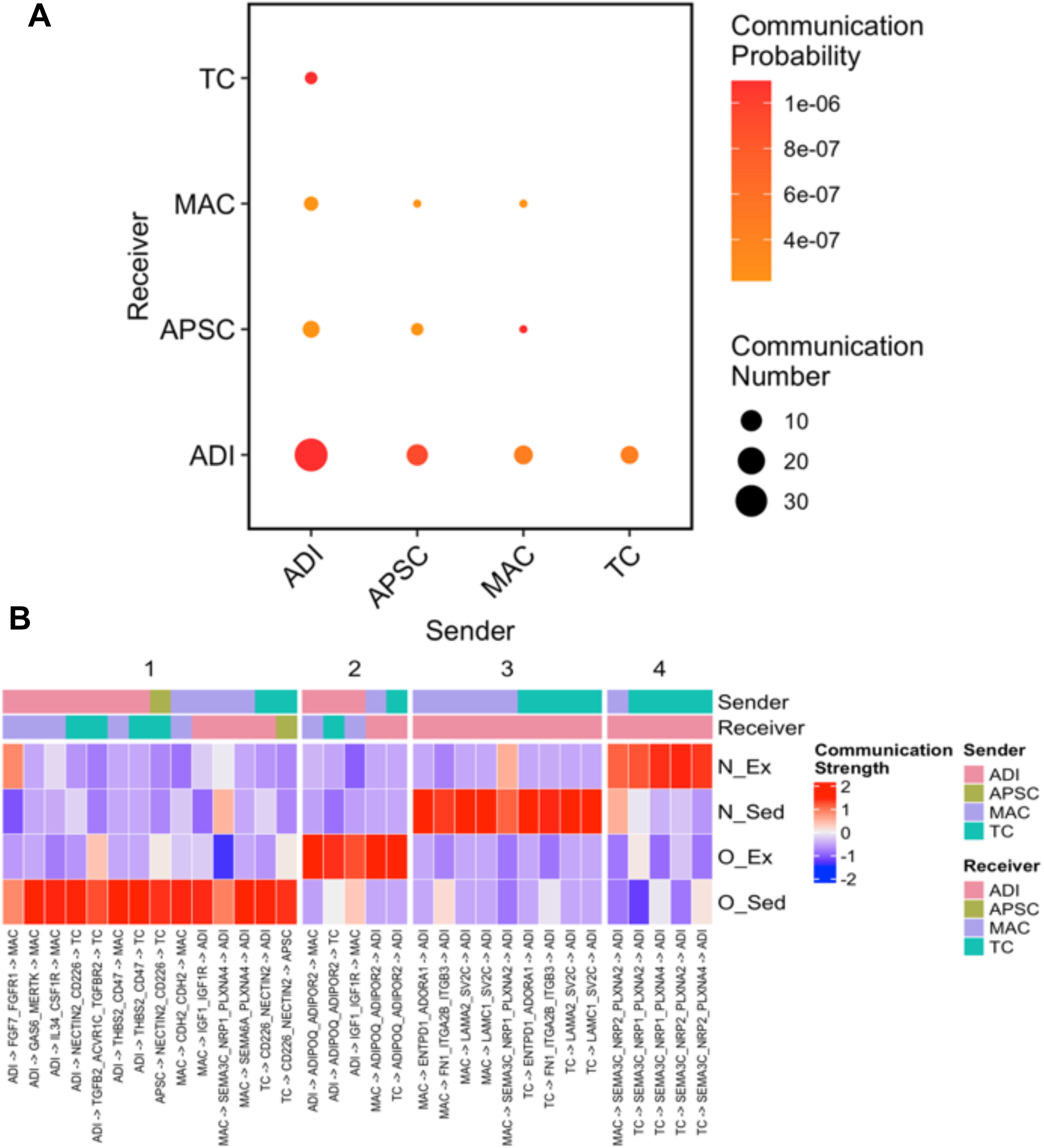
Cell-to-cell communications between immune and non-immune cells in epicardial adipose tissue. (A) Dot plot of the number of significant interactions between cell types. Circle color correlates to the probability of cell-cell interaction. Circle size correlates to the number of cell-cell interactions. (B) Heat map of the crosstalk between immune cells, adipocytes, and APSCs in EAT. Communication strength for each interaction, as well as the sender and receiver, are shown for each cell-to-cell interaction. N_Ex: occluded exercise; N_Sed: occluded sedentary O_Ex: occluded exercise; O_Sed: occluded sedentary. ADI: adipocytes. APSC: adipocyte stem cells. MAC: macrophages. TC: T cells. See the abbreviation list for gene names.

Differential communication analysis found significant alterations in the expression of numerous ligands and receptors as determined by calculated communication scores (**Figure 7B**). O_Sed EAT had the strongest cell-cell interactions (14) of any treatment group, which occurred between all four cell types examined. Adipocytes in O_Sed EAT signaled to macrophages via fibroblast growth factor 7 (*FGF7*), growth arrest-specific 6 (*GAS6*), interleukin 34 (*IL34*), and to T cells via nectin cell adhesion molecule 2 (*NECTIN2*) and transforming growth factor beta 2 (*TGFB2*). Adipocytes modulated immune cell adhesion and migration of both immune cell types via thrombospondin 2 (*THB2*) (**Figure 7B**). On the other hand, macrophages signaled to adipocytes via insulin-like growth factor 1 (*IGF1*), semaphorin 3C (*SEMA3C*), and semaphorin 6A (*SEMA6A*), which suggests invasion and adhesion of macrophages into O_Sed EAT. T cells in O_Sed EAT adhered and migrated to adipocytes and APSCs via CD226 molecule (*CD226*) to *NECTIN1* communication (**Figure 7B**). In N_Sed EAT, immune cells had nine strong signals to adipocytes without any signaling involvement of the APSCs. While macrophages in N_Sed EAT had strong signaling to adipocytes via *SEMA3C* as macrophages in O_Sed EAT did, they also had strong signals to adipocytes via *FN1*, laminin gamma subunit 1 (*LAMC1*), laminin alpha subunit 2 (*LAMA2*), and ectonucleoside triphosphate diphosphohydrolase 1 (*ENTPD1*) (**Figure 7B**). T cells in N_Sed EAT also had strong signals to adipocytes by these same factors except for *SEMA3C*.

In contrast to O_Sed EAT, O_Ex EAT did not involve APSCs in strong cell-cell signaling interactions. EAT from exercised pigs also contained many less strong cell-cell signaling interactions than EAT from sedentary pigs (10 versus 23 interactions; **Figure 7B**). Adipocytes signaled to macrophages via adiponectin, C1Q and collagen domain containing (*ADIPOQ*), and *IGF1* but signaled to T cells vis *ADIPOQ* only (**Figure 7B**). Macrophages and T cells also signaled back to adipocytes via ADIPOQ. In N_Ex EAT, macrophages and T cells had strong signaling interactions with adipocytes via *SEMA3C*.

## Discussion

Given the critical role of inflammation in the development of CAD^38^, this study was designed to build upon our initial exploration of the effects of exercise training on the cellular composition and transcript expression of EAT in female pigs with chronic ischemic heart disease.^20^ Previously, we found that exercise training increased interactions amongst immune, mesenchymal, and endothelial cells in female EAT.^20^ We applied both RNA-seq and snRNA-seq to gain additional insights into the diversity of cell types in EAT and their responses to exercise and coronary artery occlusion. Using snRNA-Seq, we clustered cell types by gene signatures. We then analyzed cell-to-cell communication between immune cells, adipocytes, and adipocyte stem cells in EAT from sedentary and exercised-trained female pigs with and without coronary artery occlusion.

Bulk RNA-seq of EAT found that exercise training increased the transcript expression of genes related to cellular growth factors and cell adhesion molecules and decreased the transcript expression of genes related to metal ion binding and transferase actions, tight junctions, and cytoskeletal structure. These results are from bulk RNA-seq, so they cannot be attributed to a particular cell type. However, genes like *CDH7,* a type II cadherin significantly upregulated in EAT from exercised pigs, are known to be responsible for slow and weak cell-to-cell adhesions^39^, as might be seen with the migration of immune cells into EAT from the periphery. Another highly upregulated growth factor in EAT from exercise-trained pigs, *HB-EGF,* is highly correlated with the movement of monocytes and dendric cells into tissues.^40^ Indeed, based on the types of highly significantly upregulated genes in EAT of exercise-trained pigs, it is not surprising that pathways enriched in this treatment group are predominantly related to the enhancement of immune cell function (i.e., chemokine signaling, phagocytosis, asthma). Interestingly, *MT2A,* which modulates intracellular zinc levels, was downregulated in EAT from exercise-trained pigs, and could be related to immune cell function in EAT. Zinc is critical for cell survival and controls apoptosis in many cell types.^41^ Moreover, zinc has been demonstrated to control T cell signal transduction^42, 43^ and regulate monocyte function^44, 45^ , possibly via nuclear factor kappa beta (NF-κβ).^46^ *CYP2B6* and *GALNTL5,* which were downregulated in EAT from exercised pigs, control metal ion binding with *CYP2B6,* specifically binding iron. Iron is increased in tissue from obese individuals, and iron overload in adipocytes has been linked to insulin resistance.^47^ Given that iron is required for adipogenesis^48^ and decreased iron in adipose tissue is related to decreased lipid uptake^49^, it makes sense that *CYP2B6* is downregulated in EAT in response to exercise. Interestingly, *CLDN4,* a tight junction regulator downregulated in the EAT from exercised pigs, is increased in visceral adipose tissue in obese individuals^50^ and has been demonstrated to be involved in T cell communication.^51^ Unlike the linear relationship between upregulated genes and enriched pathways, pathways not enriched in EAT in response to exercise don’t correlate mechanistically with the most significantly downregulated genes. Pathways that are not enriched are all related to energy production (i.e., TCA cycle, oxidative phosphorylation, pyruvate metabolism) and adipogenesis (i.e., fatty acid metabolism and PPAR signaling). The significance of this disconnect is not clear and requires further investigation.

snRNA-seq provided a more nuanced assessment of the cellular landscape of EAT, revealing unique cellular distribution patterns across each treatment group. Non-occluded EAT in both activity groups had many adipocytes, which suggests that growth factors and lipids required to support adipogenesis are likely sourced from the systemic circulation. Interestingly, APSCs were highly abundant in N_Sed EAT but not N_Ex EAT, which suggests that the N_Sed EAT has a large capacity for additional adipogenesis. Moreover, N_Sed EAT had the most T cells and macrophages of any treatment group. This finding suggests these immune cells may migrate into EAT from other locations in the body in sedentary as opposed to exercised pigs. Thus, similar to what is known to occur in visceral adipose tissue^52^, inflammation and recruitment of immune cells in EAT may occur in response to a sedentary lifestyle. Moreover, contrary to what we expected, there are more immune cells in N_Sed than O_Sed EAT, which indicates that coronary artery occlusion may prevent peripheral immune cells from trafficking to EAT.

Opposite of what was found in the analysis of the bulk RNA-seq data set, DGE analysis of individual cell types in the snRNA-seq dataset found significant upregulation in metabolic pathways in response to exercise. Irrespective of coronary artery occlusion status, *ME1* and *SCD* were upregulated in adipocytes in response to exercise. These genes are essential for the TCA cycle and fatty oxidation, indicating that exercise training increases energy production by the adipocyte pool in EAT. Adipocytes from N_Ex EAT were enriched for insulin signaling, glucose turnover pathways, and adipokine signaling. Thus, adipocytes from EAT with adequate localized blood flow likely have increased insulin sensitivity in response to exercise. Our results in EAT correspond with findings that high-intensity exercise increases visceral, but not subcutaneous, adipose tissue sensitivity.^53^ Indeed, O_Ex EAT did not have these same enriched pathways, except for PPAR signaling. Surprisingly, PPAR signaling, a master regulator of adipogenesis^54^, is enriched in adipocytes in response to exercise, irrespective of coronary artery occlusion status. This finding is at odds with human studies, which have found a reduction in EAT volume in response to endurance exercise.^55, 56^ However, PPARs in adipocytes have also been shown to inhibit inflammation-related gene transcription by blocking transcription factors such as NF𝜅𝛽, signal transducer and activator of transcription (STAT), and nuclear factor of activated T cells (NFAT).^57^ Therefore, it seems more likely that the enrichment of PPAR signaling in adipocytes in response to exercise may play an anti-inflammatory, as opposed to adipogenic, role in these cells.

In response to exercise, immune cells also had upregulation of genes related to fatty acid biosynthesis (*ACACA, ELOVL6*) and energy production (*ME1*) and enrichment of pathways such as biosynthesis of unsaturated fatty acids, pyruvate metabolism, and insulin signaling. PPAR signaling was enriched in response to exercise in all immune cells, irrespective of the occlusion status of the coronary arteries. *PPARG* activation in macrophages of mice causes suppression of immunoreactive cytokines such as tumor necrosis factor-alpha (TNF𝛼), interleukin 6 (IL6), and interleukin 1 beta (IL1𝛽)^58^ and promotion of immunotolerant factors such as interleukin 4 (IL4), CD36 molecule (CD36), and interleukin 10 (IL10).^59^ Therefore, PPARG activation in macrophages favors the transition from an M1 inflammatory to an M2 anti-inflammatory phenotype. Indeed, PPAR signaling was enriched in macrophages from exercised EAT, and fewer M1 inflammatory macrophages were found in exercised EAT. By contrast, N_Sed EAT had many M1 inflammatory macrophages compared to the other treatment groups. In contrast, O_Sed EAT had about a third of the number of M1 inflammatory macrophages that N_Sed EAT had. However, irrespective of the number of macrophages in the sedentary EAT or the coronary artery occlusion status, a significant fraction of macrophages in sedentary EAT expressed genes in the tumor necrosis factor (TNF) superfamily. TNF ligands and receptors are proinflammatory modulators of programmed cell death.^60^ Thus, from a macrophage gene expression perspective, sedentary EAT is more pro-inflammatory than exercised EAT. However, as N_Sed EAT had the largest number of M1 macrophages and increased gene expression of pro-inflammatory markers on macrophages, it is the unhealthiest EAT of all the groups.

The leukocyte transendothelial migration pathway is negatively enriched in O_Ex EAT, which supports the theory that coronary artery occlusion decreases the number of peripheral immune cells able to migrate into EAT. M0 or unpolarized macrophages are monocytes that have moved into tissue from the bloodstream and await cytokine cues to polarize into M1 or M2 macrophages.^61^ Therefore, the exceptionally high numbers of M0 macrophages in N_Sed EAT further indicate that migration of peripheral monocytes into EAT was highest in sedentary states and required intact coronary artery circulation. However, EAT from exercise-trained environments had a higher percentage of M0 macrophages than EAT from sedentary environments. Thus, exercise may dampen the release of cytokines from other immune cells that could program M0 macrophages to polarize.

Similar to our findings in macrophages, exercise training enriched PPAR signaling pathway in T cells. Most studies regarding the role of *PPARG* activation in T cells have focused on CD4+ T cells, such as T helper cells. Indeed, PPARG signals with mechanistic target of rapamycin kinase (mTOR) in CD4+ T cells to directly program fatty acid uptake to provide energy for the activation and proliferation of this T cell subpopulation.^62^ Similar to the findings in macrophages, N_Sed EAT had the highest number of T cells of any treatment group. However, unlike the findings in macrophages, O_Ex EAT also had a very high number of T cells. Indeed, UMAP analysis of O_Ex T cells demonstrated the separation of this T cell group from all other T cells in EAT. Examination of the KEGG pathway analysis provides clues regarding the possible functional differences in the T cell population from O_Ex EAT. T cells from O_Ex EAT, but not N_Ex EAT, have enrichment of pathways related to energy production and fatty acid biosynthesis, which would be required for differentiation and activation.^63^ However, T cells from O_Ex EAT have negative enrichment of T cell receptor signaling, Jak-Stat signaling, and natural killer (NK) cell mediated cytotoxicity. Jak-stat is a major regulator of T cell metabolism^64^ and cell homeostasis.^65^ T cell receptor signaling is essential for T cell activation, differentiation, and immune response.^66^ Interestingly, one of the roles of NK cells in adipose tissue is to work with T cells to secrete cytokines to polarize macrophages.^67^ Based on the functional analysis of these negatively enriched pathways, T cells from O_Ex EAT may be in an inactive, quiescent state. Many inactive T cells could be a reason for the large number of unpolarized macrophages found in O_Ex EAT. Interestingly, although CD8+ T cells were the most abundant T cell subpopulations in EAT, O_Ex and N_Ex EAT had the lowest number of CD8+ T cells. Given that O_Ex had the second largest number of T cells of any treatment group, the low number of CD8+ T cells in O_Ex EAT suggests many T cells in O_Ex EAT may be from a different T cell subpopulation, likely CD4+ T cells. In contrast, N_Sed EAT had the most T cells of any treatment group and the most CD8+ T cells of any treatment group. Therefore, most of T cells in N_Sed EAT were CD8+ T cells.

The N_Sed EAT had the largest fraction of T cells expressing genes in the TNF superfamily. O_Sed EAT had the second largest fraction of T cells expressing genes in the TNF superfamily and had a significantly higher fraction of T cells expressing TNF superfamily genes than T cells from O_Ex EAT. Therefore, T cells from N_Sed EAT were CD8+ T cells with high cytokine expression levels. Meanwhile, T cells from O_Sed EAT were likely CD8+ cells with moderate cytokine expression levels. Inflamed visceral adipose tissue, as found in obese individuals, contains activated CD8+ T cells, which secrete tumor necrosis factor-alpha (TNF𝛼), interferon gamma (INF𝛾), and tumor necrosis factor beta (TNF𝛽).^68^ Similar to what we proposed for macrophages, T cells from N_Sed EAT were likely activated CD8+ T cells from the peripheral circulation that secreted pro-inflammatory cytokines. T cells from O_Sed EAT also were likely activated CD8+ T cells with lower cytokine activity. However, further research should be done to determine the sources of these CD8+ T cells, particularly in O_Sed EAT.

Interestingly, *PPARG* was highly expressed in T cells from all treatment groups. However, when CD8+ T cells were examined, *PPARG* expression was extremely low for CD8+ T cells from N_Ex EAT. Activation of PPARG in CD8+ T cells causes a pro-inflammatory phenotype with increased production of inflammatory cytokines.^69^ Thus, not only are there very few CD8+ T cells in N_Ex EAT, their activation via PPARG is nearly nonexistent. Indeed, most T cells from N_Ex EAT could have been CD4+ T cells with moderate cytokine expression levels. By contrast, T cells from O_Ex could have been CD4+ T cells with low cytokine expression levels. T helper 1 (Th1) and Th17 cells are CD4+ T cells that play a role in tissue inflammation in obesity.^70, 71^ In contrast, T regulatory cells (Tregs) are CD4+ T cells that have an anti-inflammatory role in adipose tissue.^72, 73^ We postulate that exercise may increase the number of CD4+ T cells in the tissue and favor the Treg cell type. The next steps in this research should be to collect the occluded and non-occluded tissue from exercised and sedentary pigs to examine the concentration of cytokines in the tissue and perform flow cytometry for specific T cell subpopulations. As there were too few T cells from other subpopulations than CD8+ in our samples, future approaches will require more tissue to determine all the T cell populations in EAT.

Analysis of cell-cell communication with CellChat demonstrated that sedentary EAT had the strongest cell-cell communications. In support of our theory that N_Sed EAT had increased numbers of immune cells due to migration into the tissue from the coronary arteries, all outgoing significant strong communications in N_Sed EAT originated from macrophages or T cells to adipocytes and utilized genes related to cell adhesion and migration. Thus, immune cells could likely migrate into EAT via connections with adipocytes. O_Sed EAT had stronger cell-cell communications than N_Sed EAT. However, unlike N_Sed EAT, there was incoming and outgoing communication between immune cells and adipocytes in O_Sed EAT. Furthermore, while communication from adipocytes to immune cells utilized cell adhesion and migration genes, adipocytes also communicated to immune cells via cytokines and growth factors in O_Sed EAT. This suggests cell migration, possibly through capillaries in EAT instead of coronary arteries, and local proliferation of immune cell populations may have contributed to increased immune cells in O_Sed EAT. Interestingly, while the overall number of macrophages and T cells in O_Sed EAT was not more than the number of these cells from exercised EAT, there were more pro-inflammatory M1 macrophages and CD8+ T cells in O_Sed EAT than exercised EAT. Given that the coronary artery adjacent to the EAT was occluded, localized cell proliferation and polarization of resident cells likely occurred in O_Sed EAT. There were very few strong cell-cell communications in exercised EAT. O_Ex EAT, but not N_Ex EAT, had communication via growth factors from adipocytes to immune cells and immune cells to adipocytes. This suggests that the larger pool of CD8+ T cells and M1 macrophages found in O_Ex EAT as opposed to N_Ex EAT resulted from local tissue proliferation. However, the number of CD8+ T cells and M1 macrophages in O_Ex EAT is about half that of O_Sed EAT, suggesting the exercise mitigates the proliferation of these two pro-inflammatory immune cell classes. Under sedentary conditions, EAT releases excess low-density lipoproteins (LDL) and adipokines, such as interleukin 6 (IL6), that can elicit a pro-inflammatory response.^74, 75^ However, exercise training has been previously found to suppress visceral adipose tissue inflammation and inflammatory macrophage accumulation in obese male mice by reducing the number of M1 macrophages and CD8+ T cells.^76^ Our findings build upon what is known about the effects of exercise on visceral adipose tissue. This data underscores the importance of exercise training in the modulation of cellular communication dynamics in EAT, potentially mitigating an inflammatory status brought on by coronary artery occlusion.

In summary, our research demonstrates that a sedentary lifestyle increases the number of inflammatory M1 macrophages and CD8+ T cells, their expression of TNF superfamily genes, and the number of strong communications with adipocytes in EAT. Interestingly, the number of inflammatory M1 macrophages and CD8+ T cells in sedentary EAT is decreased when the adjacent coronary artery is occluded. However, irrespective of coronary artery occlusion status, exercise training mitigates EAT inflammation by decreasing the number of inflammatory M1 macrophages and CD8+ T cells and their expression of TNF superfamily genes. Exercise also increases the expression of the anti-inflammatory factor *PPARG* by adipocytes and macrophages which likely mitigates the number of M1 macrophages in exercised EAT. These findings emphasize that exercise training, even in the presence of coronary artery occlusion, creates an anti-inflammatory milieu in EAT. Future studies will focus on how the anti-inflammatory milieu of EAT in the exercised female communicates with and modulates the function of the surrounding cardiac structures.

## Acknowledgments

We thank Jeff Bray and Andrew Hillhouse for their assistance with this work.

## Supplemental Material

Supplemental Datasets (Excel format)

Supplemental Figures and Figure Legends (PDF file format)

Supplemental Tables and supporting information (PDF file format)

## Abbreviations and Acronyms

ACACA: acetyl coA carboxylase alpha
ACLY: ATP citrate lyase
ACTA2: actin alpha 2, smooth muscle
ADIPOQ: adiponectin, C1Q and collagen domain containing
ARHGAP28: Rho GTPase activating protein 28
CAD: coronary artery disease
CCL5: C-C motif chemokine ligand 5
CD163: CD163 molecule
CD2: CD2 molecule
CD8A: CD8A molecule
CD80: CD80 molecule
CD209: CD209 molecule
CDH7: cadherin 7
CDH13: cadherin 13
COL1A2: collagen type alpha 2 chain
CRIM1: cysteine rich transmembrane BMP regulator 1
CSF1R: colony stimulating factor 1 receptor
DAB2: DAB adaptor protein 2
DEG: differentially expressed genes
DGAT2: diacylglycerol O-acyltransferase 2
EAT: epicardial adipose tissue
EGFR: epithelial growth factor receptor
ELOVL6: ELOVL fatty acid elongase 6
FABP5: fatty acid binding protein 5
FASN: fatty acid synthase
FGF7: fibroblast growth factor 7
FN1: fibronectin 1
GALNT16: polypeptide N-acetyl galatosaminyltransferase 16
GAS6: growth arrest-specific 6
GJC1: gap junction protein gamma 1
GMZH: granzyme H
GPM6A: glycoprotein M6A
GPX3: glutathione peroxidase 3
GZMH: granzyme H
IGF1: insulin-like growth factor 1
INF𝛾: interferon gamma
IKZF3: IKAROS family zinc finger 3
IL34: interleukin 34
IL6: interleukin 6
IRF5: interferon regulatory factor 5
KEGG: Kyoto encyclopedia of genes and genomes
LAMA2: laminin subunit alpha 2
LDB2: LIM domain binding 2
LDL: low density lipoprotein
LPL: lipoprotein lipase
ME1: malic enzyme 1
MRVI1: inositol 1,4,5-triphosphate receptor associated 1
mTOR: mechanistic target of rapamycin kinase
NCALD: neurocalcin delta
NECTIN1: nectin cell adhesion molecule 2
NFAT: nuclear factor of activated T cells
NK: natural killer
NF𝜅𝛽: nuclear factor kappa beta
NRXN1: neurexin 1
PCA: principal component analysis
PDGFRA: platelet-derived growth factor receptor alpha
PDK4: pyruvate dehydrogenase kinase 4
PICK1: protein interacting with PRKCA 1
PKHD1L1: PKHD1 Like 1
PLIN: perilipin 1
POSTN: periostin
PPAR: peroxisome proliferator-activated receptor
PPARD: peroxisome proliferator-activated receptor delta
PPARG: peroxisome proliferator-activated receptor gamma
RBPJ: recombination signaling binding protein for immunoglobulin kappa J region
SCARA5: scavenger receptor class A member 5
SCD: stearoyl-coA desaturase
SEMA3C: semaphorin 3C
SEMA6A: semaphorin 6A
SKAP1: Src kinase-associated phosphoprotein 1
SNCA: synuclein alpha
STAB1: stabilin 1
STAT: signal transducer and activator of transcription
SYNPO2: synaptopodin 2
TGFB1: transforming growth factor beta 1
TGFB2: transforming growth factor beta 2
TGFBR2: transforming growth factor receptor 2
THB2: thrombospondin 2
TNF: tumor necrosis factor
TNF𝛼: tumor necrosis factor alpha
TNF𝛽: tumor necrosis factor beta
TOX: thymocyte selection associated high mobility group box
UMAP: uniform manifold approximation and projection
UMI: unique molecular identifier
VWF: von Willebrand factor

