## Supplemental Figures for "Aerobic exercise decreases the number and transcript expression of inflammatory M1 macrophages and CD8+ T cells in the epicardial adipose tissue of female pigs"

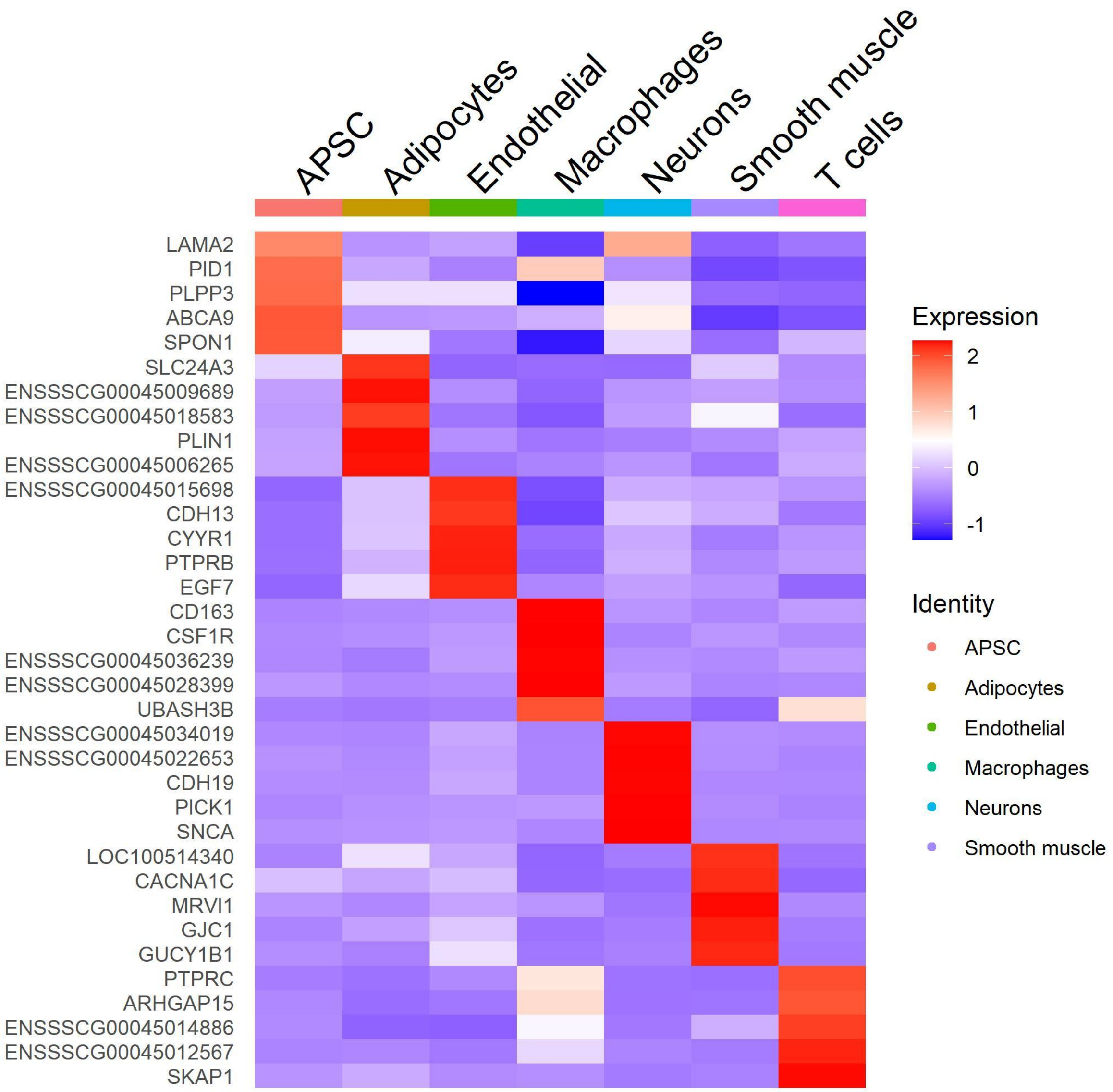


**Figure S1: Expression of top transcript markers for each cell type found in single nuclei sequencing of epicardial adipose tissue from exercise-trained and sedentary female pigs.** Red is upregulated, and magenta is downregulated expression level of the top 35 genes expressed in all cell types. As this study was conducted in porcine epicardial adipose tissue, many of the genes have not been identified by homologous name and function to human genes and, as such, are shown by their Ensembl Stable Id. Refer to Supplemental Dataset S2.

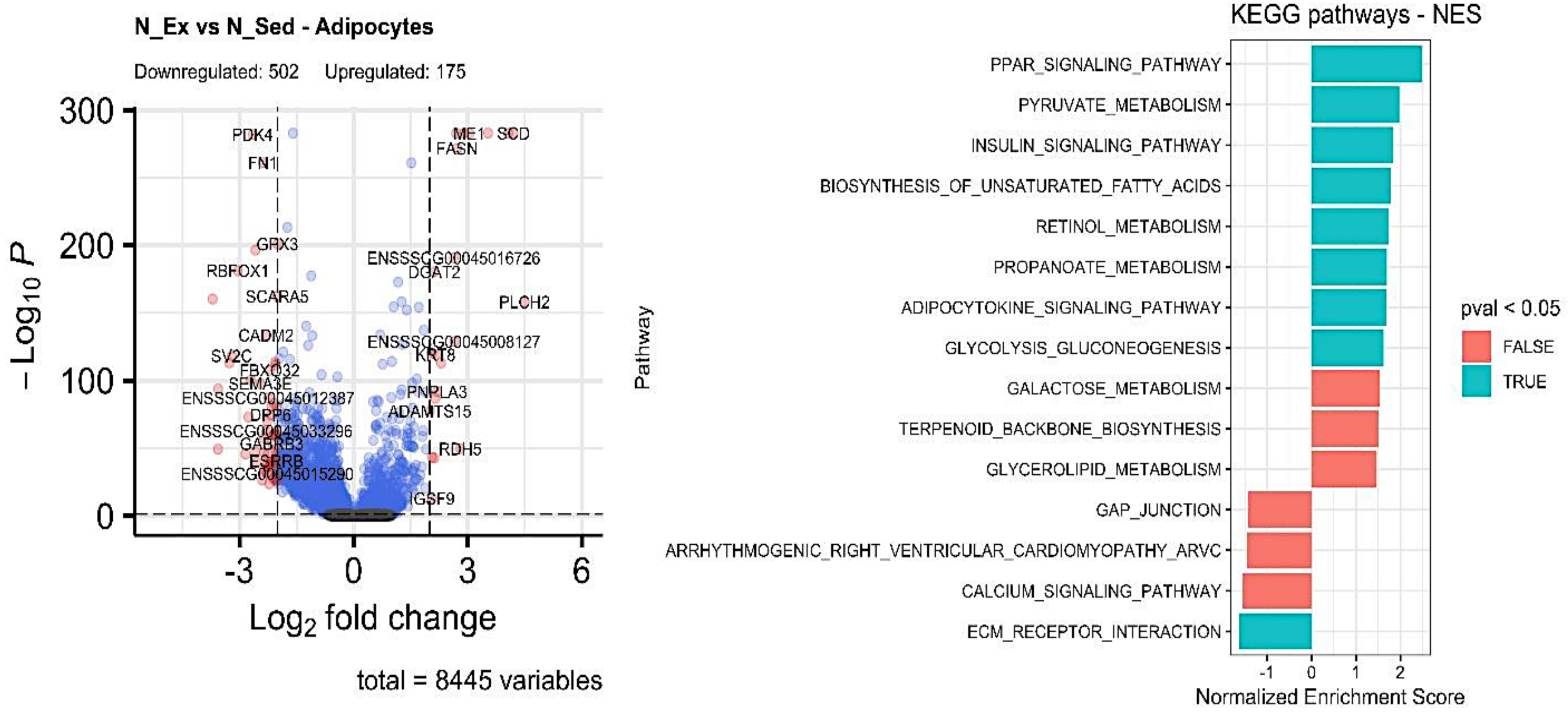


**Figure S2. Analysis of differentially expressed genes within adipocytes in non-occluded epicardial adipose tissue from exercised and sedentary female pigs.** Visual representation of the volcano map depicting upregulated and downregulated genes. The threshold set points for DEG were $\log_2$ fold change > 2 and adjusted P value < 0.05. For the volcano map, if a gene was significantly upregulated or downregulated, it is shown in red. If a gene was significantly regulated but did not have a $\log_2$ fold change > 2, it is shown in blue. Identification of KEGG enriched (positive NES) and non-enriched (negative NES) pathways in exercise-trained pigs. Teal blue: significant; Orange; non-significant. P < 0.05. N_Ex: non-occluded exercise; N_Sed: non-occluded sedentary. Refer to Supplemental Dataset S3.

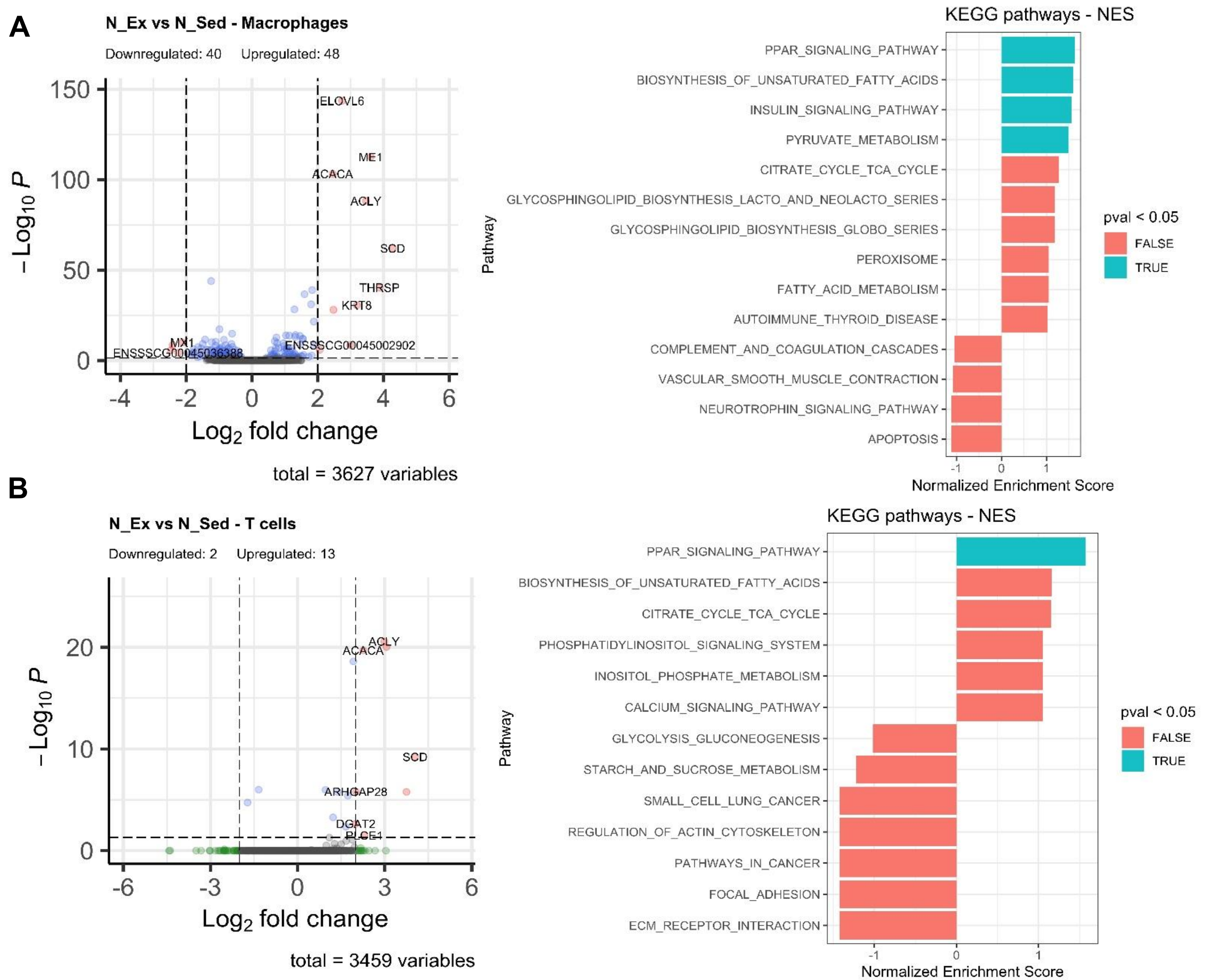


**Figure S3. Analysis of differentially expressed genes within macrophages (A) and T cells (B) in non-occluded epicardial adipose tissue from exercised and sedentary female pigs.** Visual representation of the volcano map depicting upregulated and downregulated genes. The threshold set points for DEG were $\log_2$ fold change > 2 and adjusted P value < 0.05. For the volcano map, if a gene was significantly upregulated or downregulated, it is shown in red. If a gene was significantly regulated but did not have a $\log_2$ fold change > 2, it is shown in blue. Identification of KEGG enriched (positive NES) and non-enriched (negative NES) pathways in exercise-trained pigs. Teal blue: significant; Orange; non-significant. $P < 0.05$. N_Ex: non-occluded exercise; N_Sed: non-occluded sedentary. Refer to Supplemental Dataset S3.

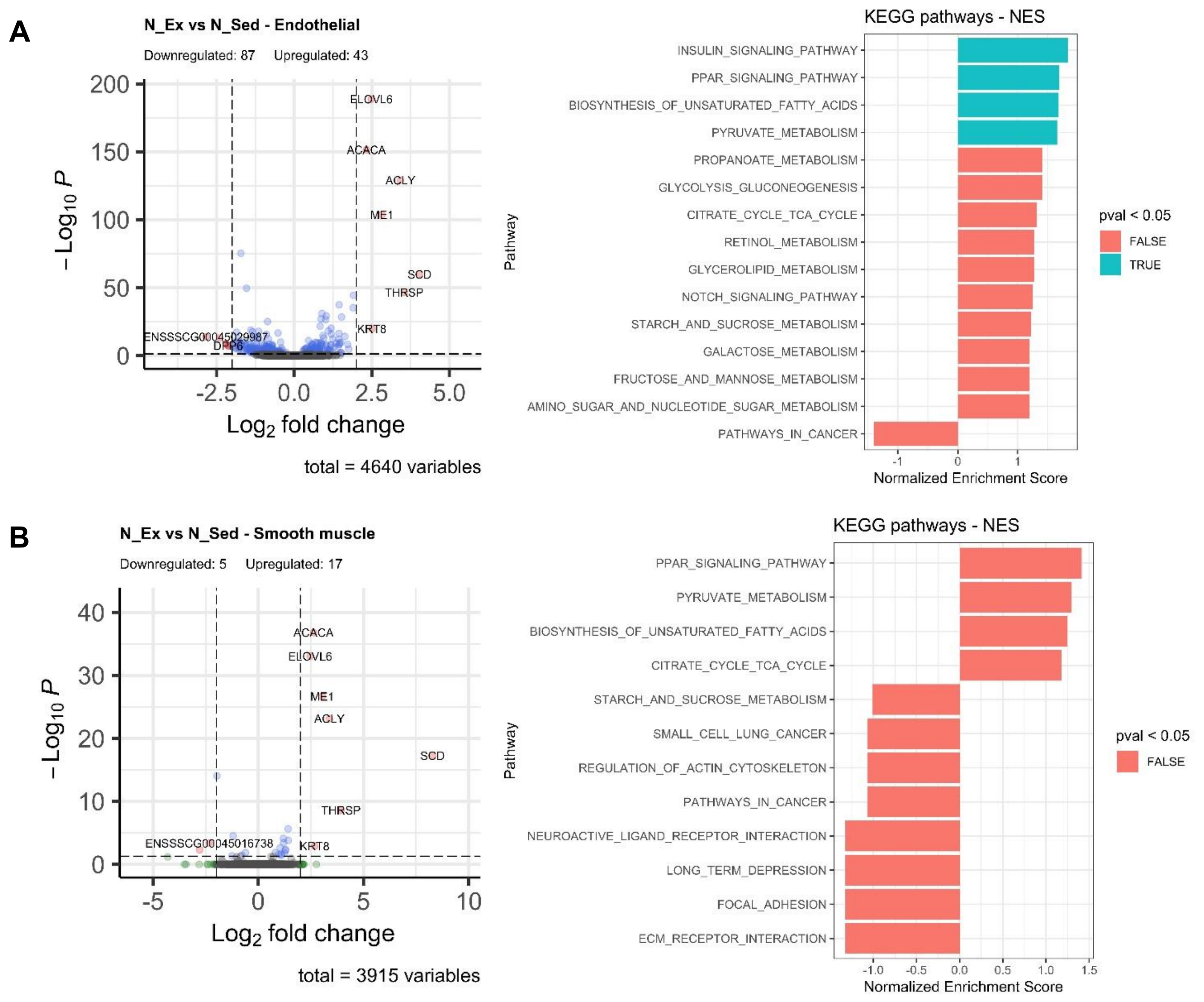


**Figure S4. Analysis of differentially expressed genes within endothelial (A) and smooth muscle cells (B) in non-occluded epicardial adipose tissue from exercised and sedentary female pigs.** Visual representation of the volcano map depicting upregulated and downregulated genes. The threshold set points for DEG were $\log_2$ fold change > 2 and adjusted P value < 0.05. For the volcano map, if a gene was significantly upregulated or downregulated, it is shown in red. If a gene was significantly regulated but did not have a $\log_2$ fold change > 2, it is shown in blue. Identification of KEGG enriched (positive NES) and non-enriched (negative NES) pathways in exercise-trained pigs. Teal blue: significant; Orange; non-significant. P < 0.05. N_Ex: non-occluded exercise; N_Sed: non-occluded sedentary. Refer to Supplemental Dataset S3.

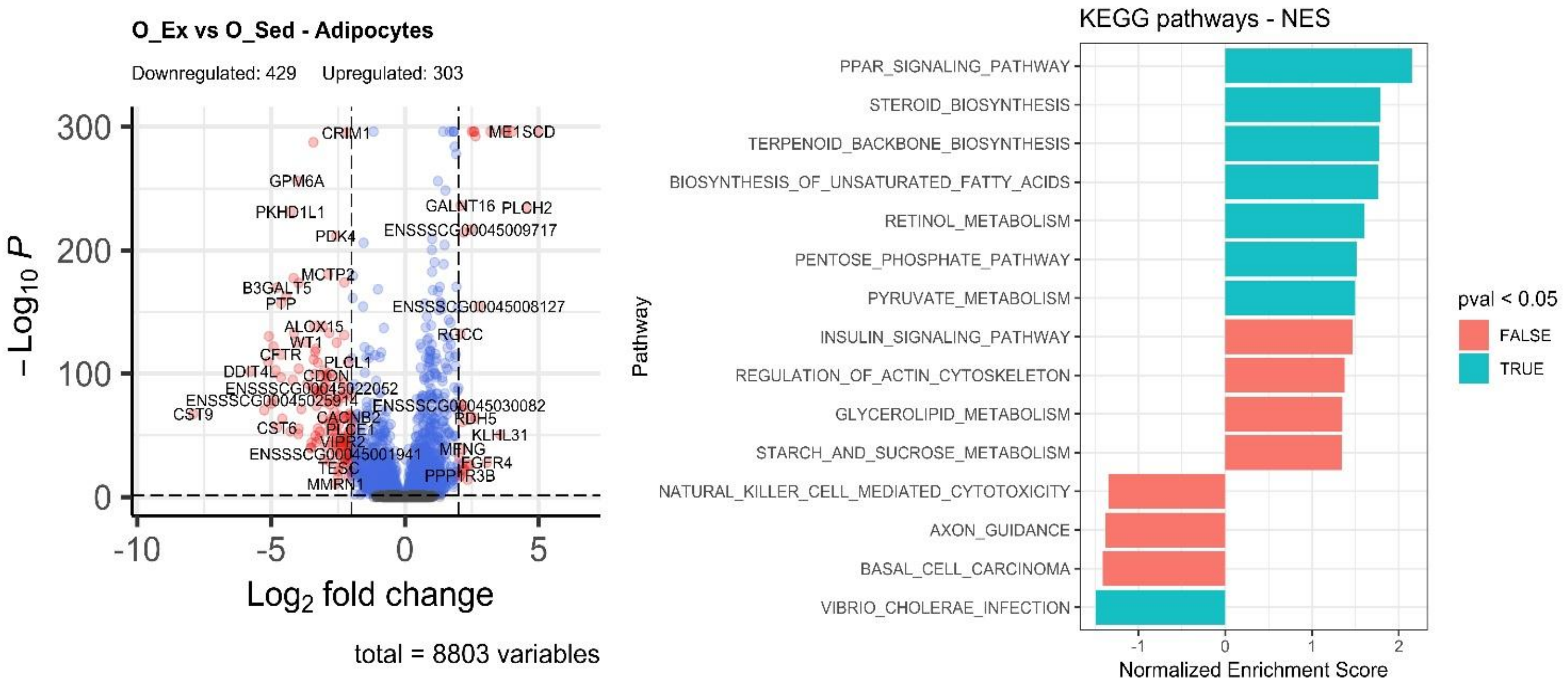


**Figure S5. Analysis of differentially expressed genes within adipocytes in occluded epicardial adipose tissue from exercised and sedentary female pigs.** Visual representation of the volcano map depicting upregulated and downregulated genes. The threshold set points for DEG were $\log_2$ fold change > 2 and adjusted P value < 0.05. For the volcano map, if a gene was significantly upregulated or downregulated, it is shown in red. If a gene was significantly regulated but did not have a $\log_2$ fold change > 2, it is shown in blue. Identification of KEGG enriched (positive NES) and non-enriched (negative NES) pathways in exercise-trained pigs. Teal blue: significant; Orange; non-significant. P < 0.05. O_Ex: occluded exercise; O_Sed: occluded sedentary. Refer to Supplemental Dataset S4.

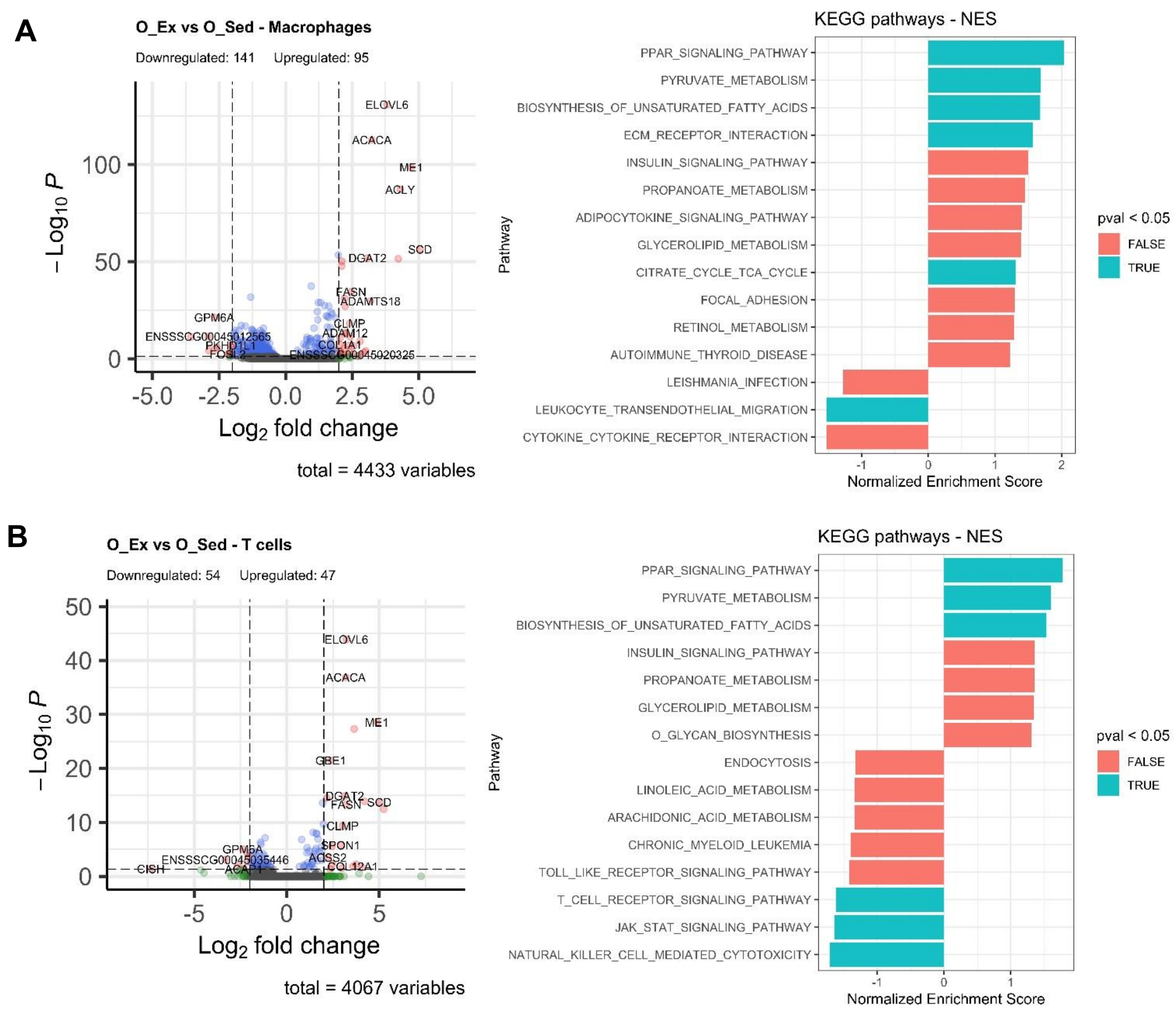


**Figure S6. Analysis of differentially expressed genes within macrophages (A) and T cells (B) in occluded epicardial adipose tissue from exercised and sedentary female pigs.** Visual representation of the volcano map depicting upregulated and downregulated genes. The threshold set points for DEG were $\log_2$ fold change > 2 and adjusted P value < 0.05. For the volcano map, if a gene was significantly upregulated or downregulated, it is shown in red. If a gene was significantly regulated but did not have a $\log_2$ fold change > 2, it is shown in blue. Identification of KEGG enriched (positive NES) and non-enriched (negative NES) pathways in exercise-trained pigs. Teal blue: significant; Orange; non-significant. P < 0.05. O_Ex: occluded exercise; O_Sed: occluded sedentary. Refer to Supplemental Dataset S4.

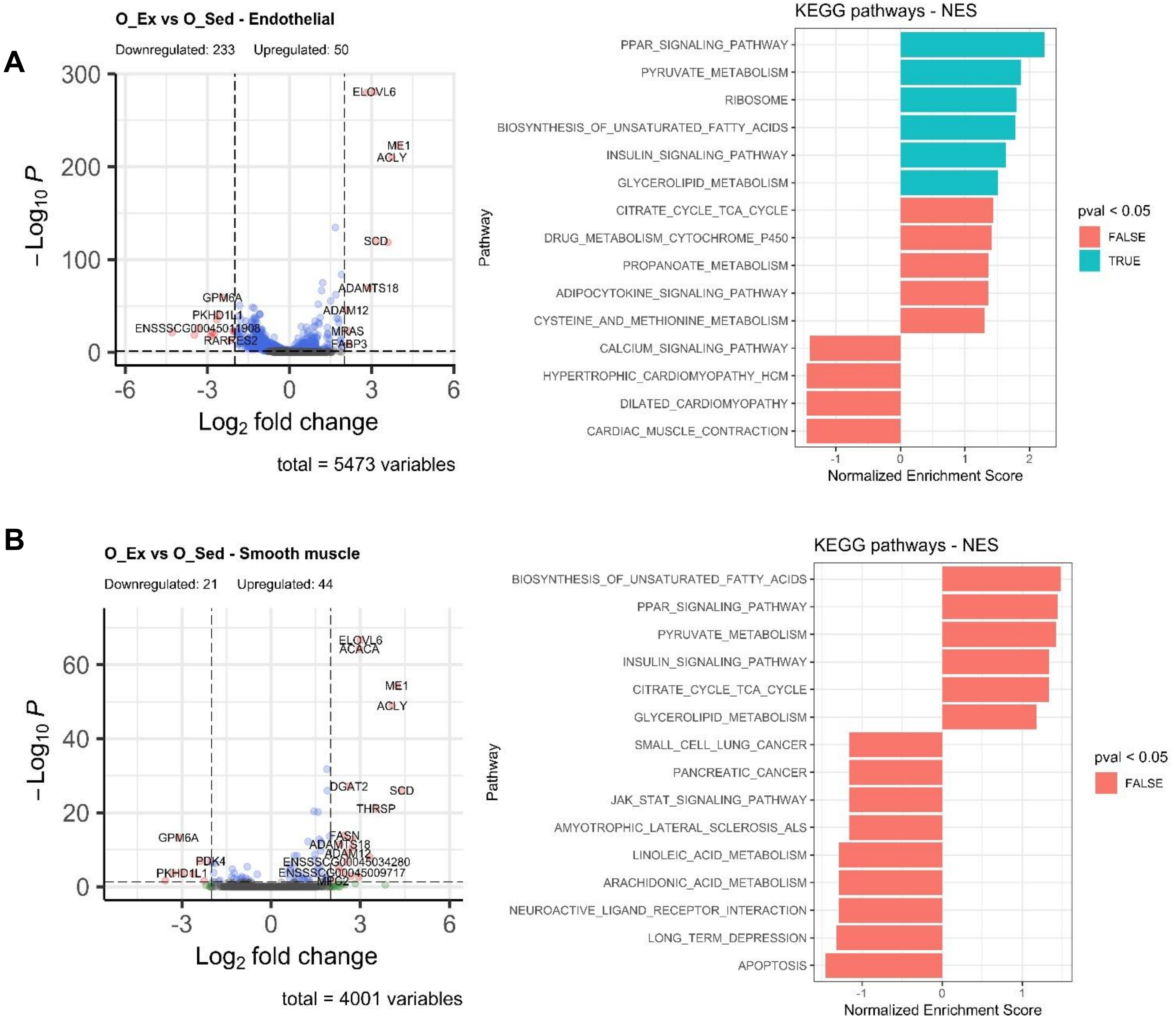


**Figure S7. Analysis of differentially expressed genes within endothelial and smooth muscle cells in occluded epicardial adipose tissue from exercised and sedentary female pigs.** Visual representation of the volcano map depicting upregulated and downregulated genes. The threshold set points for DEG were $log_2$ fold change > 2 and adjusted P value < 0.05. For the volcano map, if a gene was significantly upregulated or downregulated, it is shown in red. If a gene was significantly regulated but did not have a $log_2$ fold change > 2, it is shown in blue. Identification of KEGG enriched (positive NES) and non-enriched (negative NES) pathways in exercise-trained pigs. Teal blue: significant; Orange; non-significant. $P < 0.05$. O_Ex: occluded exercise; O_Sed: occluded sedentary. Refer to Supplemental Dataset S4.

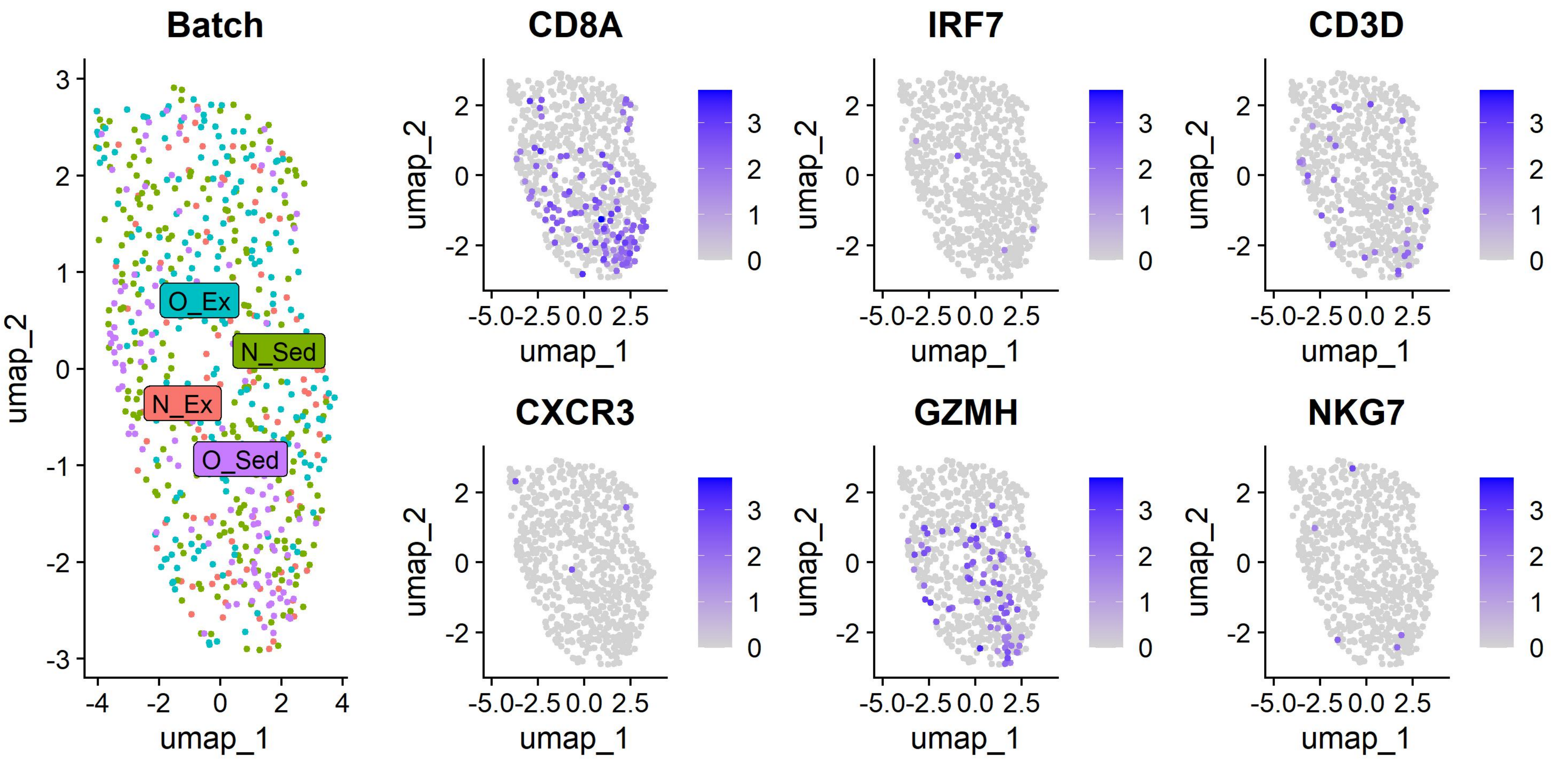


**Figure S8. Batch analysis of T cells in epicardial adipose tissue from female pigs.** No T cell separation by treatment group was seen in the UMAP. UMAP analysis of markers for subpopulations of T cells revealed many CD8+ T cells as determined by the expression of granzyme H (*GZMH*) and CD8A molecule (*CD8A*). Other genes evaluated included: interferon regulatory factor 7 (*IRF7*), CD3D molecule (*CD3D*), C-X-C Motif Chemokine Receptor 3 (*CXCR3*), and Natural Killer Cell Granule Protein 7 (*NKG7*). N_Ex: occluded exercise; N_Sed: occluded sedentary O_Ex: occluded exercise; O_Sed: occluded sedentary.

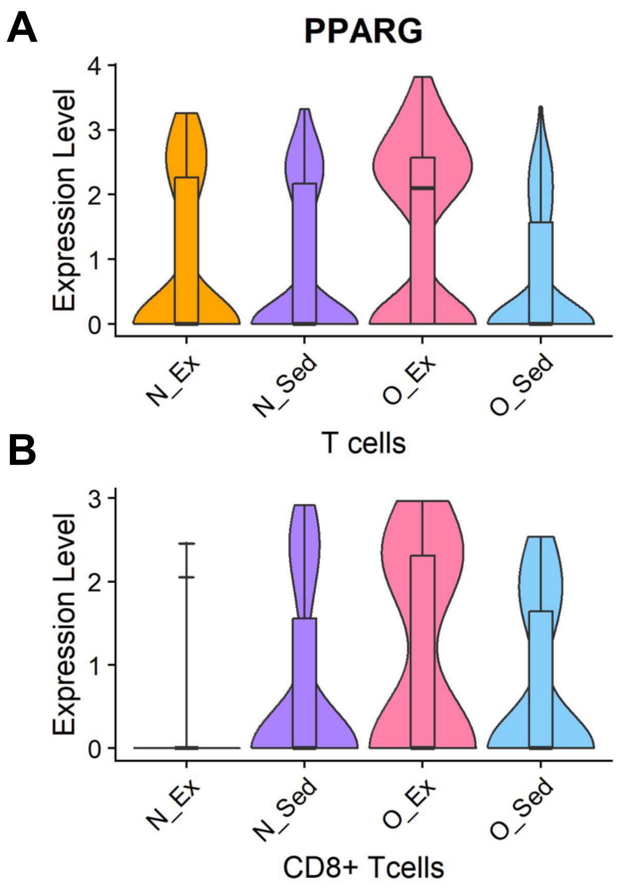


**Figure S9. PPARG expression in all T cells (A) and in CD8+ T cells (B) in epicardial adipose tissue from female pigs.** N_Ex: occluded exercise; N_Sed: occluded sedentary O_Ex: occluded exercise; O_Sed: occluded sedentary.

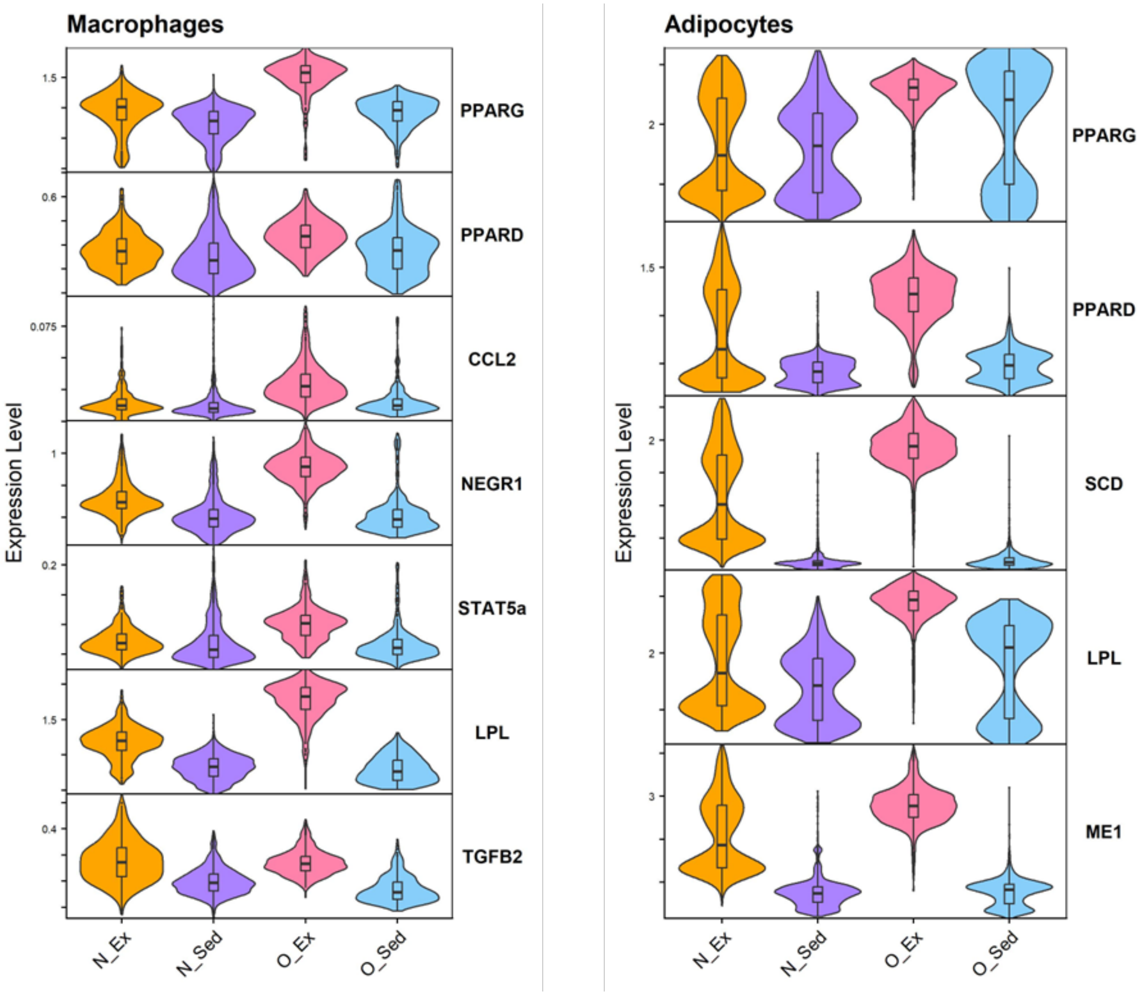


**Figure S10. The expression level of selected marker genes involved in PPARG signaling in macrophages and adipocytes from epicardial adipose tissue.** N_Ex: occluded exercise; N_Sed: occluded sedentary O_Ex: occluded exercise; O_Sed: occluded sedentary.

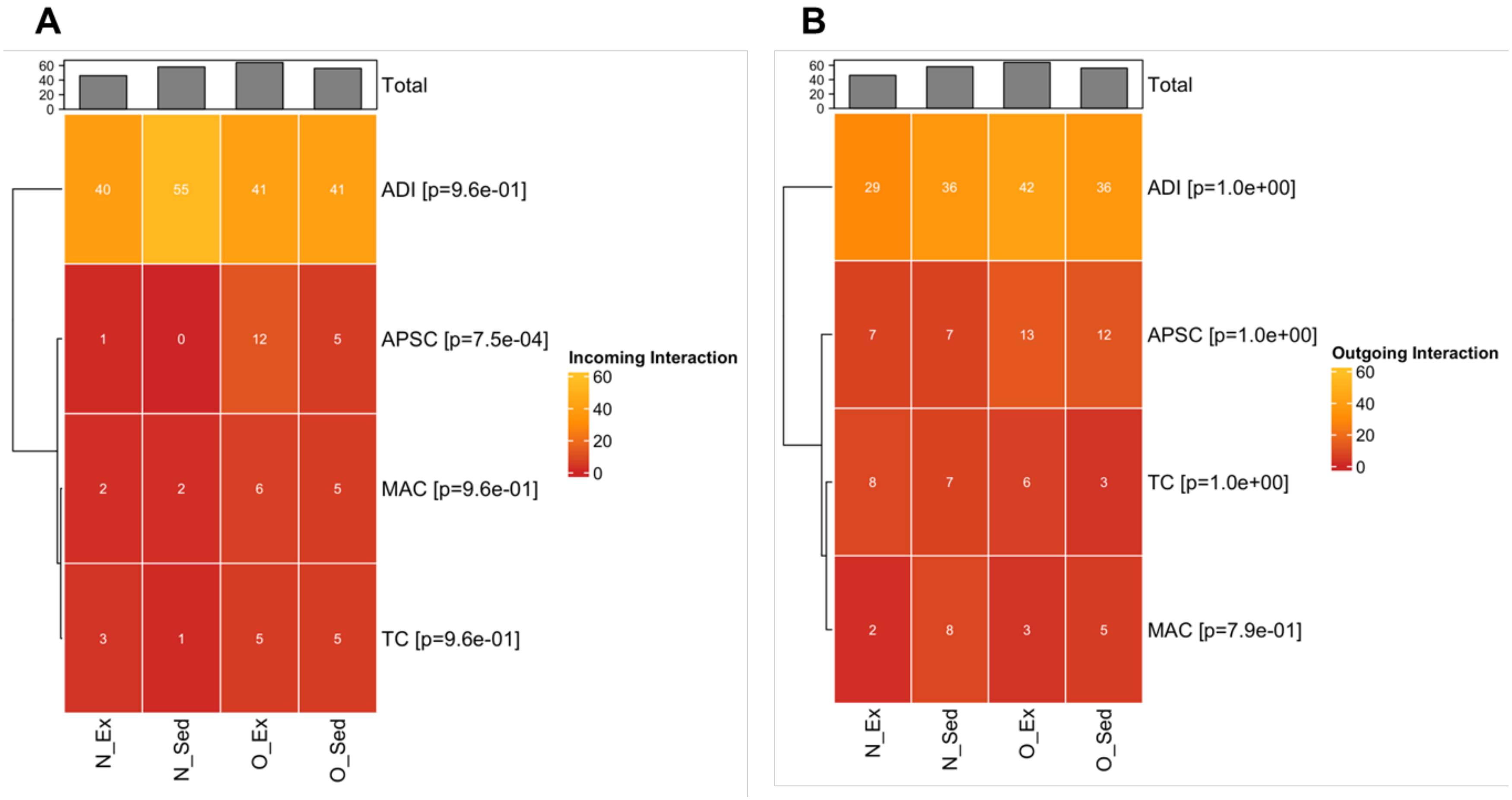


**Figure S11**. **Cell-to-cell interaction number between four cell types found in epicardial adipose tissue**. Heatmaps show the total number of significant incoming (A) and outgoing (B) interactions for each cell type**.** N_Ex: occluded exercise; N_Sed: occluded sedentary O_Ex: occluded exercise; O_Sed: occluded sedentary. ADI: adipocytes. APSC: adipocyte stem cells. MAC: macrophages. TC: T cells.
