## Supplemental Tables for "Aerobic exercise decreases the number and transcript expression of inflammatory M1 macrophages and CD8+ T cells in the epicardial adipose tissue of female pigs"

**Table S1. Transcript markers for each cell type identified in epicardial adipose tissue from female pigs**

| Cell Type | Transcript Markers |
|---|---|
| Adipocytes | PLIN1, ADIPOQ, DGAT2, PKHD1L1, ME1 |
| Adipocyte progenitor stem cells (APCS) | PDGFRA, EGFR, SCARA5, LAMA2, COL1A2, POSTN |
| Macrophages | STAB1, CD163, RBPJ, DAB2, CSF1R |
| T cells | NCALD, SKAP1, TOX, IKZF3 |
| Endothelial Cells | VWF, FABP5, CDH13, LDB2 |
| Smooth muscle cells | ACTA2, GJC1, SYNPO2, MRVI1 |
| Neurons | SNCA, NRXN1, PICK1 |

**Table S2. Numbers of cells in each treatment group by cell type**

| | Cell Type | | | | | | |
|---|---|---|---|---|---|---|---|
| **Treatment Group** | Adipocytes | APSC | Endothelial | Macrophages | Neurons | Smooth muscle | T cells |
| N_Ex | 3437 | 1795 | 820 | 417 | 72 | 167 | 104 |
| N_Sed | 2985 | 2371 | 1042 | 1199 | 100 | 242 | 254 |
| O_Ex | 2389 | 2689 | 1049 | 417 | 68 | 238 | 202 |
| O_Sed | 1793 | 2031 | 1100 | 419 | 60 | 246 | 122 |

N_Ex: non-occluded exercised; N_Sed: non-occluded sedentary. O_Ex: occluded exercised; O_Sed: occluded sedentary; APSC: adipocyte stem cells
